# SubClonal Hierarchy Inference from Somatic Mutations: automatic reconstruction of cancer evolutionary trees from multi-region next generation sequencing

**DOI:** 10.1101/011833

**Authors:** Noushin Niknafs, Violeta Beleva-Guthrie, Daniel Q. Naiman, Rachel Karchin

**Affiliations:** Department of Biomedical Engineering and Institute for Computational Medicine, Johns Hopkins University, Baltimore, MD, USA; Department of Applied Math and Statistics, Johns Hopkins University, Baltimore, MD, USA; Department of Oncology, Johns Hopkins Medical Institutions, Baltimore, MD, USA

## Abstract

Recent improvements in next-generation sequencing of tumor samples and the ability to identify somatic mutations at low allelic fractions have opened the way for new approaches to model the evolution of individual cancers. The power and utility of these models is increased when tumor samples from multiple sites are sequenced. Temporal ordering of the samples may provide insight into the etiology of both primary and metastatic lesions and rationalizations for tumor recurrence and therapeutic failures. Additional insights may be provided by temporal ordering of evolving subclones – cellular subpopulations with unique mutational profiles. Current methods for subclone hierarchy inference tightly couple the problem of temporal ordering with that of estimating the fraction of cancer cells harboring each mutation. We present a new framework that includes a rigorous statistical hypothesis test and a collection of tools that make it possible to decouple these problems, which we believe will enable substantial progress in the field of subclone hierarchy inference. The methods presented here can be flexibly combined with methods developed by others addressing either of these problems. We provide tools to interpret hypothesis test results, which inform phylogenetic tree construction, and we introduce the first genetic algorithm designed for this purpose. The utility of our framework is systematically demonstrated in simulations. For most tested combinations of tumor purity, sequencing coverage, and tree complexity, good power (≥ 0.8) can be achieved and Type 1 error is well controlled when at least three tumor samples are available from a patient. Using data from three published multi-region tumor sequencing studies of (murine) small cell lung cancer, acute myeloid leukemia, and chronic lymphocytic leukemia, in which the authors reconstructed subclonal phylogenetic trees by manual expert curation, we show how different configurations of our tools can identify either a single tree in agreement with the authors, or a small set of trees, which include the authors’ preferred tree. Our results have implications for improved modeling of tumor evolution and the importance of multi-region tumor sequencing.

**Author Summary:** Cancer is a genetic disease, driven by DNA mutations. Each tumor is composed of millions of cells with differing genetic profiles that compete with each other for resources in a process similar to Darwinian evolution. We describe a computational framework to model tumor evolution on the cellular level, using next-generation sequencing. The framework is the first to apply a rigorous statistical hypothesis test designed to inform a new search algorithm. Both the test and the algorithm are based on evolutionary principles. The utility of the framework is shown in computer simulations and by automated reconstruction of the cellular evolution underlying murine small cell lung cancers, acute myeloid leukemias and chronic lymophocytic leukemias, from three recent published studies.

## Introduction

The clonal evolution hypothesis in cancer states that cancer genomes are shaped by numerous rounds of cellular diversification, selection and clonal expansion [1, 2]. Recent methods to characterize tumor clonal evolution can be divided into two broad classes – sample tree reconstruction and subclone tree reconstruction. The first class of methods models the history of clonal evolution in an individual as a phylogenetic tree with leaves being the individual’s tumor samples, yielding a relative temporal ordering and estimate of divergence between the samples [3, 4, 5]. The second class aims at reconstructing the history of clonal evolution as a tree, which summarizes lineage relationships between cellular subpopulations [6, 7, 8, 9].

Until single-cell sequencing data is widely available, accurate high resolution modeling of tumor evolution [10] will likely remain exceedingly difficult, if not impossible. On the other hand, the coverage depth of current next generation sequencing experiments limits the number of cellular subpopulations or *subclones* detectable in tumor samples to a few (approximately 5-10) that have undergone signficant clonal expansions[11, 4, 12, 13, 14]. Each of these subclones emerge from a parental population of cells by acquiring additional somatic mutations, and cells within each subclone can be assumed to be homogeneous. Modeling of subclone evolution often involves estimating the fraction of cancer cells harboring each somatic mutation *i.e.*, somatic mutation *cellularity*, which can be inferred from next generation sequencing read count data. For example, PyClone [15] employs a Markov Chain Monte Carlo method to identify groups of mutations with similar cellularities, and SciClone [16] uses variational Bayes mixture models to cluster somatic mutations by their read count frequencies, which can be a proxy for cellularities.

Most recently, methods that couple the problems of somatic mutation clustering and phylogenetic reconstruction have emerged. PhyloSub applies a tree-structured stick breaking process that introduces tree-compatible cellularity values for mutation clusters [6]. A combinatoric approach based on an approximation algorithm for binary tree paritions [7] and a mixture deconvolution algorithm [8] have also been developed. However to our knowledge, most recently published studies of multi-region tumor sequencing continue to employ manual curation to construct a subclone phylogeny, after mutation cellularity has been estimated computationally [12, 13, 14].

We propose that progress in methods to reconstruct subclonal phylogenies will be substantially enabled by decoupling the problems of temporal ordering of subclones from that of mutation cellularity estimation. The SubClonal Hierarchy Inference from Somatic Mutations (SCHISM) framework described here can incorporate a variety of methods to estimate the cellularity of individual mutations, the cellularity of mutation clusters, and to build phylogenetic trees. First, we derive a novel mathematical formulation of assumptions about lineage precedence and lineage divergence in tumor evolution that have been fundamental to other subclone tree reconstruction methods [8, 6, 7, 9]. Lineage precedence is modeled in terms of a statistical hypothesis test, based on a generalized likelihood ratio. Hypothesis test results are combined with lineage divergence assumptions and formulated as a fitness function that can be used to rank tree topologies, generated by a phylogenetic algorithm. In this work, we designed an implementation of genetic algorithms to build phylogenetic trees. However, the fitness function can also be combined with other approaches to phylogenetic tree reconstruction. The hypothesis test can be combined with any method to estimate mutation or cluster cellularities to infer their temporal orderings.

We use simulations to evaluate the power of the hypothesis test and show that for many combinations of tumor purity, sequencing coverage, and phylogenetic tree complexity, the hypothesis test has good power (≥ 0.8) and Type 1 error is well controlled, when at least three samples from a patient are available. The simulations also confirm that the problem of subclonal phylogenetic tree reconstruction is underdetermined in many settings when the tumor sample count per individual is smaller than the number of subclones *i.e*, nodes in the phylogenetic tree. In these cases, we may see that the genetic algorithm identifies multiple equally plausible phylogenetic trees. However, when the problem is sufficiently determined, in general when the number of samples equals or exceeds the tumor sample count, the genetic algorithm reliably reconstructs the true tree, in most combinations listed above.

Using data from three published multi-region tumor sequencing studies of murine small cell lung cancer [13], acute myeloid leukemia [12] and chronic lymophocytic leukemia [14], we show how SCHISM can be configured with a variety of inputs. For all samples in these three studies, SCHISM identified either a single tree in agreement with the tree reconstructed manually by the authors, or a small set of trees, which include the authors’ published tree.

## Results

### Simulations

#### Generalized likelihood ratio hypothesis test

The hypothesis test yielded good power on average and Type 1 error was well controlled (Fig. 1). Power improved as the number of samples per individual increased. As the number of nodes in the subclone tree increased, yielding a more complex tree, more samples were required to achieve the same level of power. Even at the lowest purity level (50%) included in our experiments, good power (*≥* 0.8) was achieved with 1000X coverage and three or more samples.

**Figure 1.**
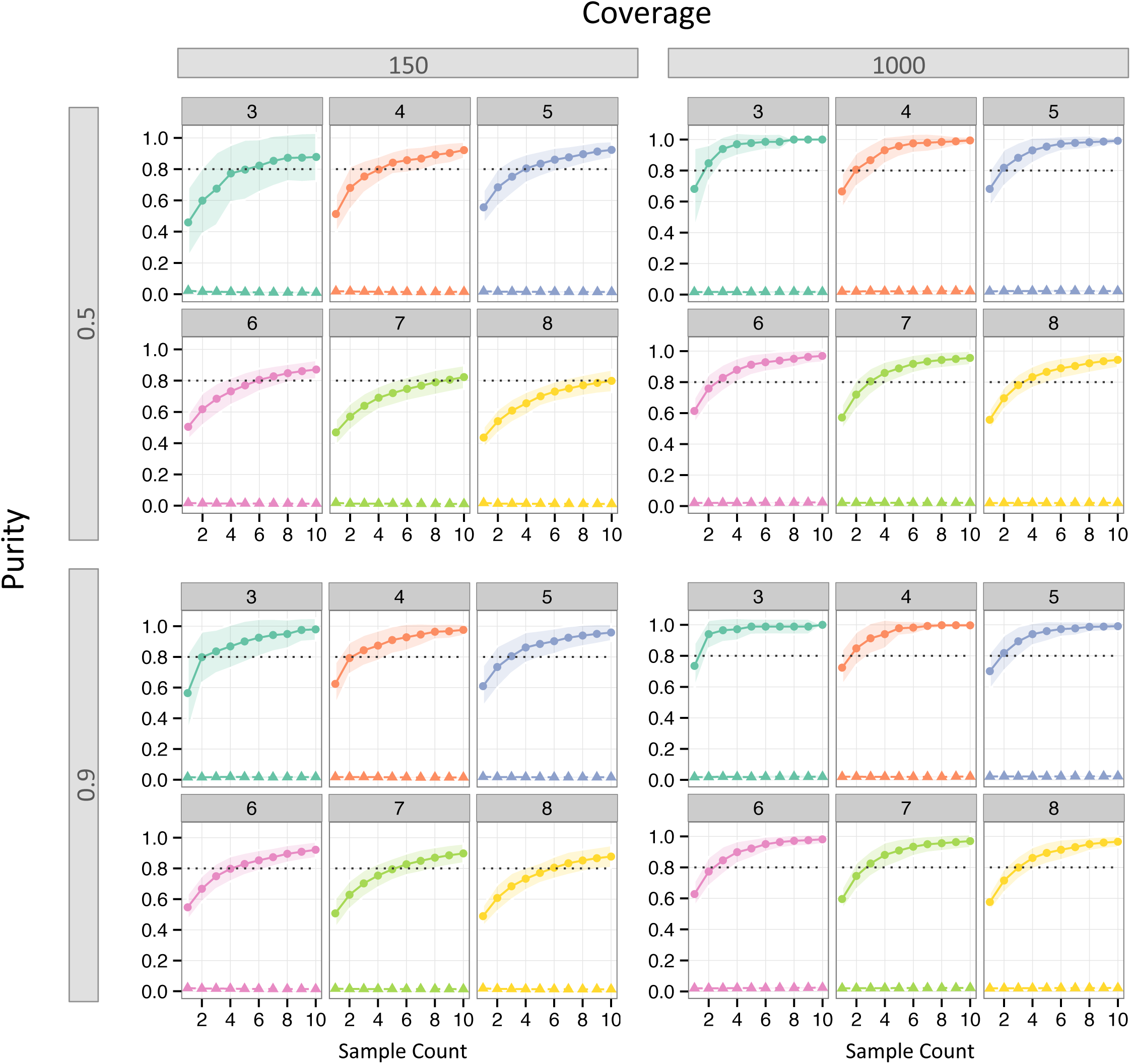
Power and Type 1 error of hypothesis test on simulated data. For each combination of coverage and purity, results are shown for trees with node counts from three to eight. Each curve was computed by taking the mean over all *instances* and all replicates for each node count. Curves with circular marks show power and curves with triangular marks show Type 1 error. Transparent coloring indicates *±*1*SE*. Dotted line indicates power=0.8.

#### Automated subclone tree reconstruction

The performance of the genetic algorithm (GA) used for tree reconstruction varied substantially, depending on the simulation inputs: number of tumor samples, node count and topology of the true tree, mutation cluster cellularity values, tumor purity, and sequencing coverage (Fig. 2). Given sample count exceeding or equal to node count, the GA most frequently (with probability ≥ 0.5) identified the true tree or a pair of maximum fitness trees that included the true tree. This probability increased to ≥ 0.75 at high purity (0.9) and coverage (1000X). As expected, simpler trees, *e.g.*, 3- or 4-node trees, were frequently identified even when the sample count was small. As trees grew more complex, a larger sample count was required, and even the most complex trees in the simulation, which had 8 nodes, were identified frequently when 10 samples were available. However, we also identified combinations of inputs for which the GA had limited success in finding the true tree. We decomposed the performance of the GA into two stages. In Stage 1, we assessed whether the tree reconstruction problem was sufficiently determined by our inputs, meaning that a single maximum fitness tree or a pair of two maximum fitness trees was identified. The GA was more likely to fail in Stage 1 when sample count was smaller than node count (Fig. 2 A1,B1). Furthermore, the settings of purity and sequencing coverage used in our simulations had less of an effect on Stage 1 success than sample count and tree node count. In Stage 2, we assessed whether the single maximum fitness or pair of maximum fitness trees included the true tree. Samples counts ≥ 5 had the most stable Stage 2 success rates, and the correct tree was recovered with increasing frequency, given higher sample counts, coverage, and purity. As expected, probability of Stage 2 success was higher for trees with smaller node counts. Our estimates of Stage 2 success were noisy when sample count was small and node count high. This behavior was a result of higher failure rates at Stage 1 under these conditions (Fig. 2 A2,B2).

**Figure 2.**
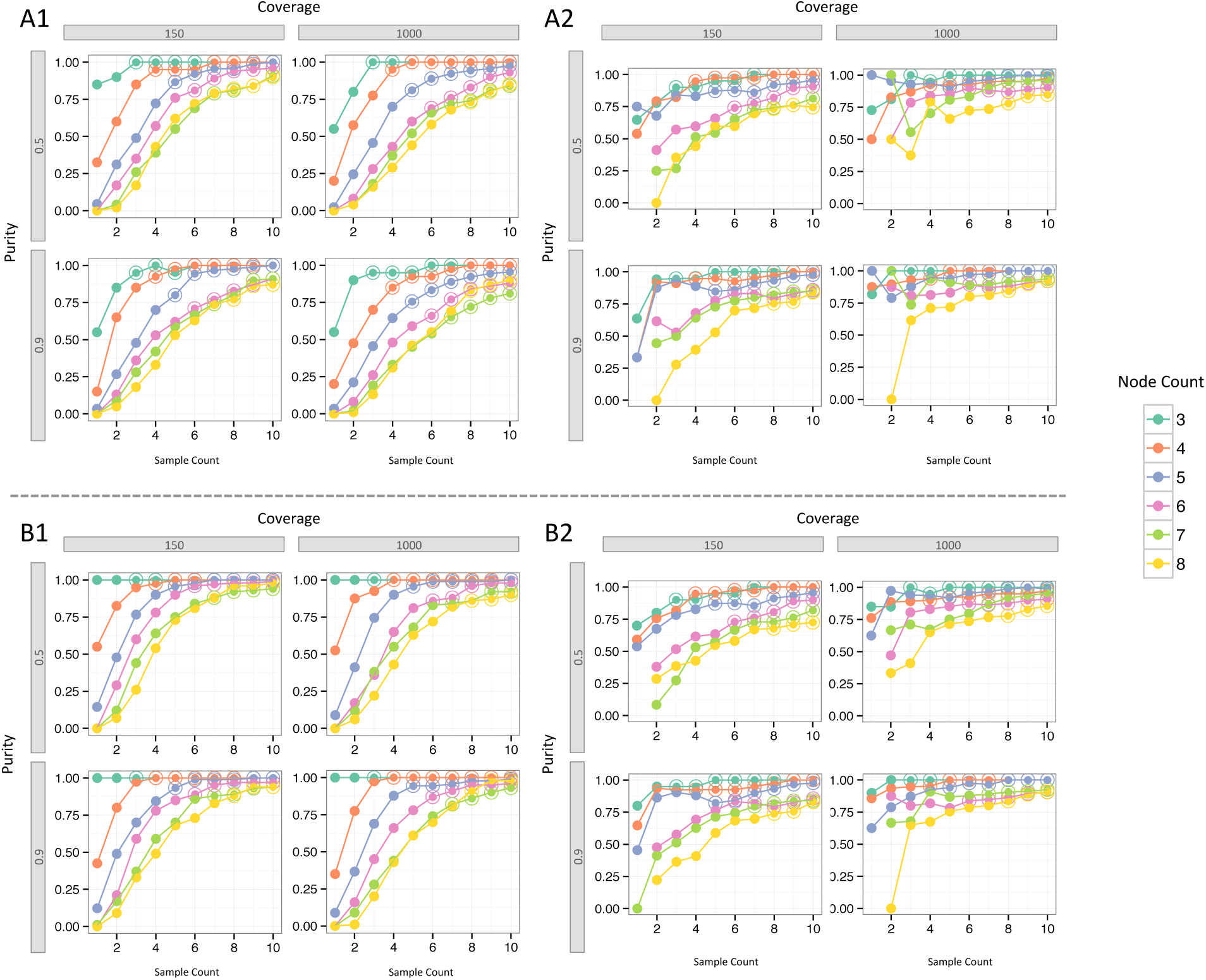
Performance of the genetic algorithm evaluated in two stages. Stage 1: Fraction of simulation runs where the genetic algorithm’s fitness function identified either a single maximum fitness tree (A1) or two maximum fitness trees (B1). Stage 2: Given success in stage 1, fraction of simulation runs where the correct tree was either the single maximum fitness tree (A2) or one of the top two maximum fitness trees (B2). For each combination of coverage and purity, results are shown for trees with node counts from three to eight. Simulations where (sample count) ≥ (node count) are marked by a double circle.

### Multi-sample sequencing studies

Recent studies of small cell lung cancer (SCLC) in mice [13], acute myeloid leukemia (AML) [12], and chronic lymphocytic leukemia (CLL) [14] attempted to infer the subclonal phylogeny underlying tumor progression, based on sequencing of multiple tumor samples. All of the studies applied computational methods to cluster somatic mutations. In some cases mutation cluster cellularities were provided, while in others read counts or cluster mean variant allele fraction was provided. The authors did not use computational methods to reconstruct the subclonal phylogenies.

We applied SCHISM to these datasets, using a variety of configurations. In cases where mutation cluster cellularities were available [13], we used the hypothesis test on pairs of mutation clusters. If mean variant allele fraction for clusters was available [12], we inferred cellularity as described in (Eq. S4) and used the hypothesis test on pairs of mutation clusters. When only read counts and mutation cluster assignments were available [12, 14], we used our own naive estimator to derive cellularity values, and applied the hypothesis test to pairs of mutations. For all configurations, we constructed a precedence order violation matrix for all pairs of mutations (*POV matrix*) or all pairs of mutation clusters (*CPOV matrix*) and ran the genetic algorithm. This approach consistently identified phylogenies identical to those manually constructed by the authors as either the single maximum fitness tree or among a small set trees tied for the maximum fitness, in underdetermined cases. In the AML study, the authors did not construct phylogenies for three out of eight patients, but they predicted which of two general clonal evolution models best explained relapse in these three patients. For two of these patients, our phylogenies were in agreement with the authors’ clonal evolution model. For the third patient, our phylogenies suggested that either of the two clonal evolution models might explain relapse. Methods details are all described in Methods (sections on Hypothesis test, Precedence Order Violation Matrix, Application to mutation clusters, Vote Aggregation and Subclonse size estimation) and Supplementary Methods (sections on Naive mutation cellularity estimate and Cluster cellularity estimation).

#### Murine small cell lung carcinoma

This study sequenced small cell lung cancer (SCLC) tumor samples from a cohort of transgenic mice with lung-specific Trp53 and Rb1 compound deletion. The full study included whole exome sequencing (WES) (150X coverage) of 27 primary tumors and metastases from six individual animals [13]. The authors applied ABSOLUTE to call copy number alterations, combine them with mutation read counts, and estimate cellularity values of each mutation. Mutation clusters were defined by clustering mutations with similar cellularity values. For three of these animals (with identifiers 3588, 3151, 984), the authors manually built subclonal phylogenetic trees.

Using the hypothesis test on the ABSOLUTE cluster mean cellularities (Fig. 5 in [13]), we generated a cluster-level precedence order violation matrix (CPOV) and a cluster cellularity by sample matrix for each mouse.

Animals 3588 and 3151 had data available for one primary and two metatastic tumors. For animal 3588, SCHISM identified an 8-node single maximum fitness tree and that tree (Fig. 3A) was identical to the authors’ manually built tree. For animal 3151, SCHISM identified six 9-node maximum fitness trees, and one of these trees was identical to the authors’ tree. While the problem was insufficiently determined, interestingly the trees shared a significant number of lineage relationships, and the main discrepancies among trees were the parental lineage for Clone3 and Clone2b (using notation from [13]) (Fig. 3B). For animal 984, data was available for one primary one metastatic tumor. SCHISM identified six 7-node maximum fitness trees (*i.e.,* underdetermined problem), and one of these trees was identical to the authors’ tree (Fig. 3C).

**Figure 3.**
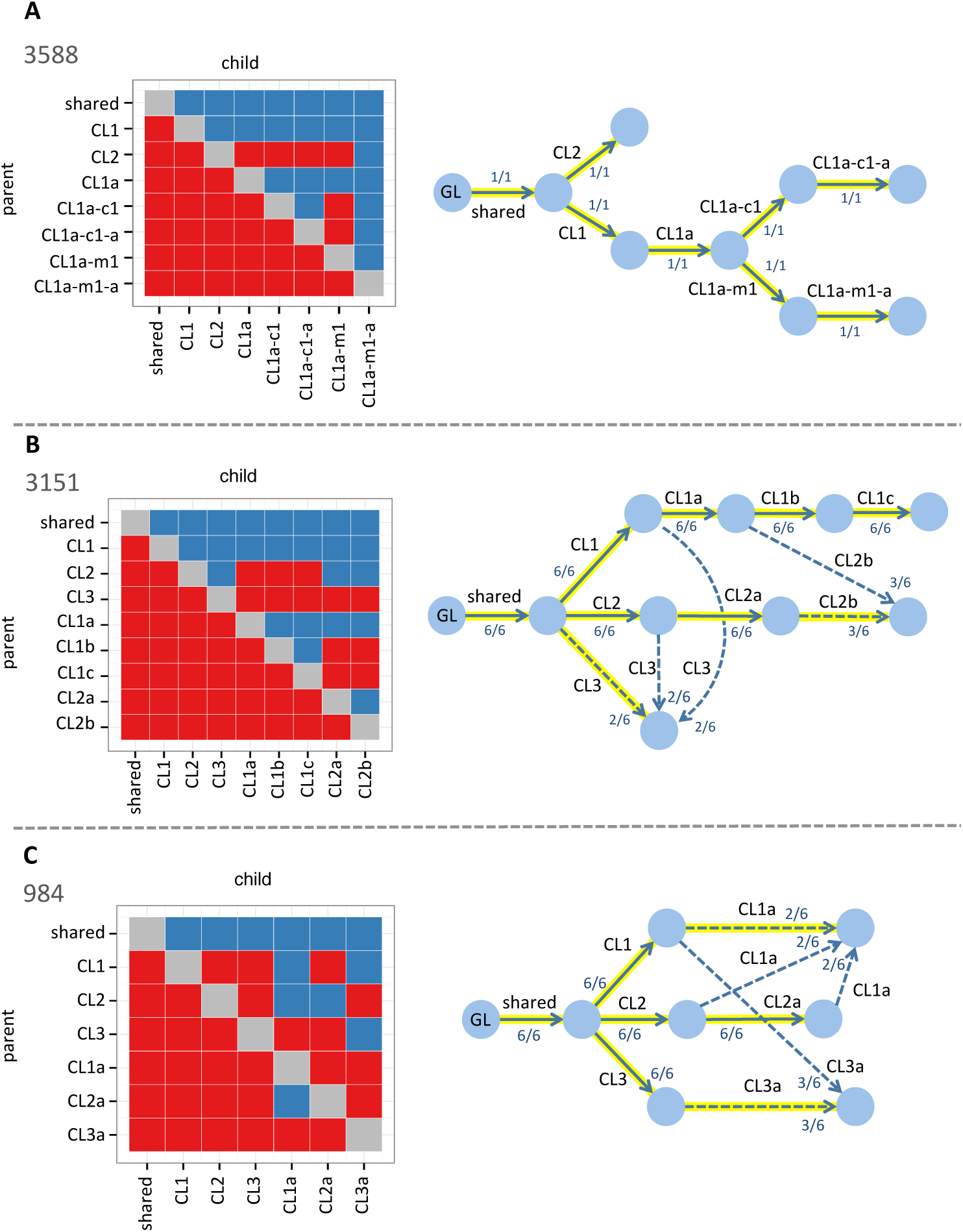
Reconstruction of subclonal phylogenies in murine models of SCLC. **A. Animal 3588.** SCHISM identified a single maximum fitness 8-node tree using one primary and two metastatic tumors. **B. Animal 3151.** Six maximum fitness 9-node trees were identified using one primary and two metastatic tumors. **C. Animal 984.** Six maximum fitness 7-node trees were identified using one primary and one metastatic tumor. Solid arrows represent lineage relationships shared by all six trees and dashed arrows represent lineage relationships shared by only a subset of the trees. Each arrow is labeled with the fraction of maximum fitness trees that include the lineage relationship. Highlighted arrows indicate the tree manually constructed by the study authors. GL = germline state. Cluster precedence order violation (CPOV) matrices are shown to the left of each tree. Columns and rows represent subclones (or Clones in the terminology of [13]). Each red square represents a pair of subclones (I,J) for which the null hypothesis that I could be the parent of J was rejected. Each blue square represents a pair for which the null hypothesis could not be rejected.

#### Chronic lymphocytic leukemia

This longitudinal study of subclonal evolution in B-cell CLL tracked three patients (CLL 003, CLL006, CLL077) over a period of up to seven years [14]. For each patient, five longitudinal peripheral blood samples were collected, and each sample was whole-genome sequenced. For selected somatically mutated sites, they further applied targeted deep sequencing (at reported 100,000X coverage). K-means clustering and expert curation were used to infer mutation clusters and subclonal phylogenetic trees were manually built.

We used mutation cluster assignments and read counts from targeted deep sequencing, for each mutation in each sample (Tables S6, S7, or S8 in [14]). Purity was estimated by identifying the mutation cluster with the maximum mean variant allele frequency in each sample. Next, based on purity and read count, we calculated cellularity (and standard error) for each mutation in diploid or copy number=1 regions with the naive estimator. The hypothesis test was performed for each pair of mutations and a cluster precedence order violation (CPOV) matrix was constructed, using a vote aggregation scheme (Methods).

For patients CLL003 and CLL077, SCHISM identified a single 4-node maximum fitness tree that was identical to the authors’ manually built tree (Fig. 4A,C). For patient CLL006, two 5-node maximum fitness trees were identified, and one was identical to the authors’ tree (Fig. 4B).

**Figure 4.**
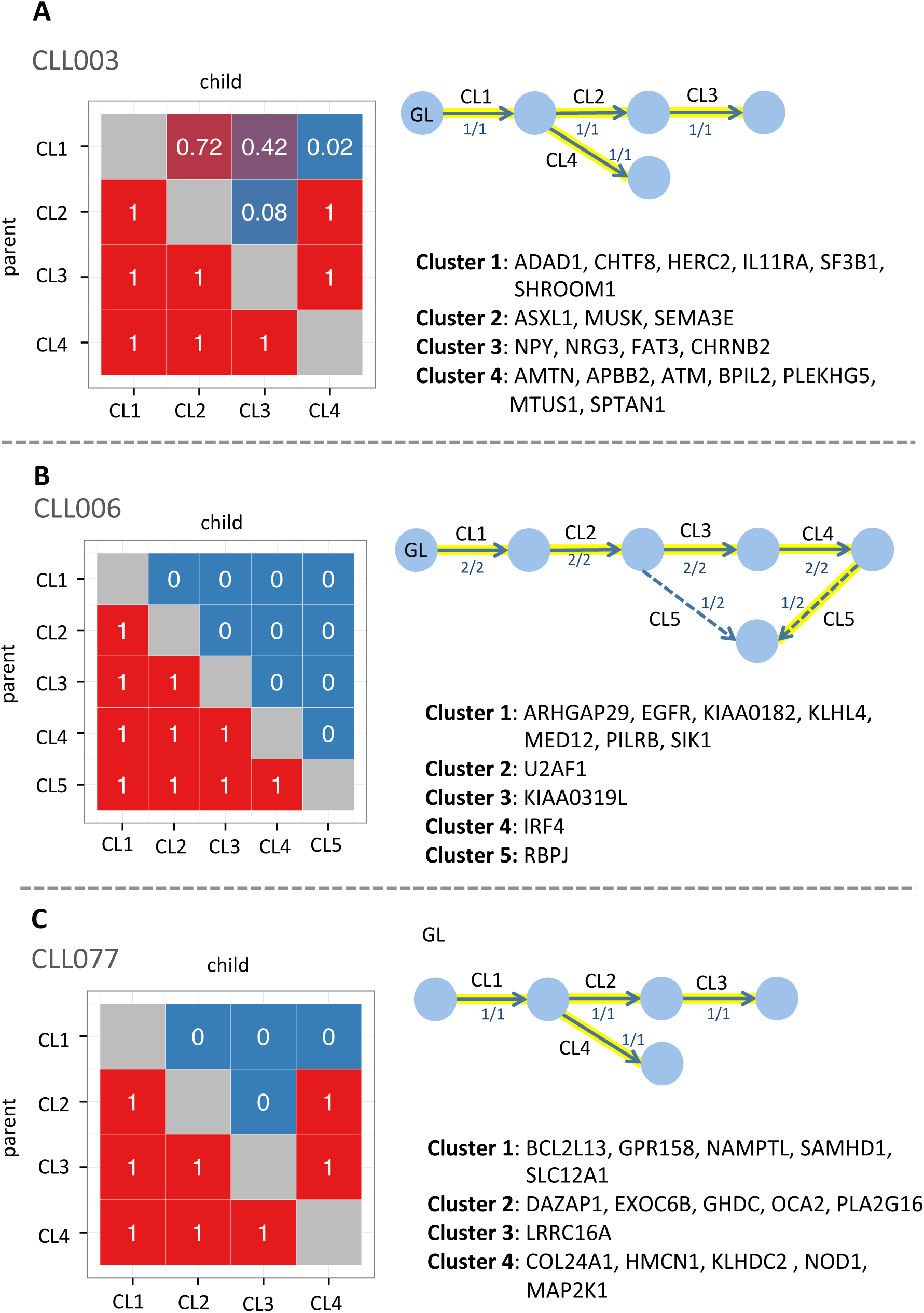
Reconstruction of subclonal phylogenies in CLL. **A. CLL003.** SCHISM identified a single maximum fitness 4-node tree using 5 samples. **B. CLL006.** Two maximum fitness 5-node trees were identified using 5 samples. Solid arrows represent lineage relationships shared by both trees and dashed arrows represent lineage relationships specific to one of the trees. **C. CLL077.** A single maximum fitness 4-node tree was identified using 5 samples. Each arrow is labeled with the fraction of maximum fitness trees that include the lineage relationship. Highlighted arrows indicate the tree manually constructed by the study authors. GL = germline state. Cluster precedence order violation (CPOV) matrices are shown to the left of each tree. Columns and rows represent mutation clusters. Each square represents a pair of mutation clusters (I,J) and the numeric value in the square shows the fraction of mutation pairs (i,j) for which the null hypothesis was rejected (Section Vote Aggregation). The mutated genes assigned to each cluster in [14, 6] are listed.

**Figure 5.**
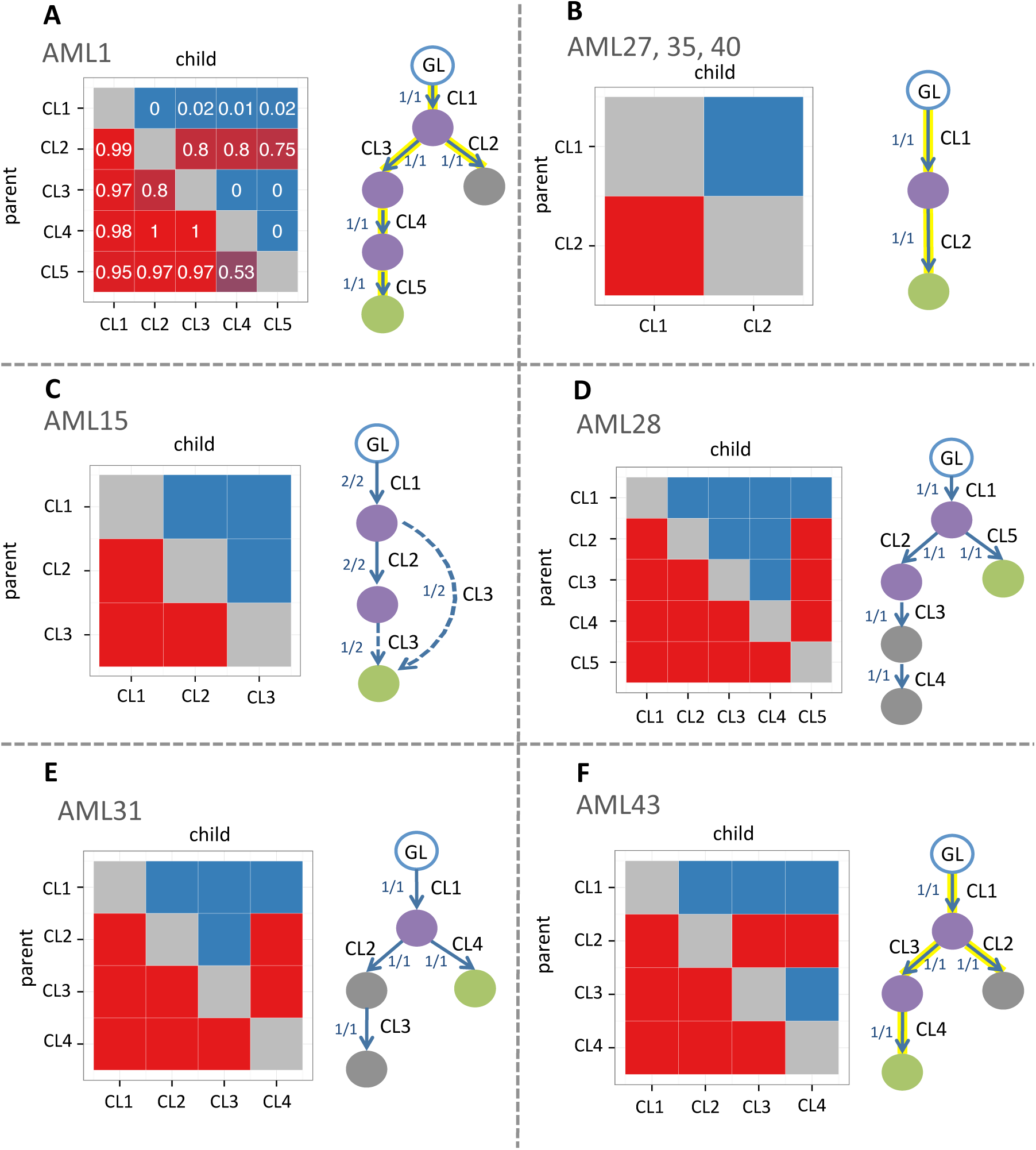
Reconstruction of subclonal phylogenies in AML. For each patient, two samples were available from primary and relapse cancers. Purple nodes represent mutation clusters present in both primary and relapse samples. Gray nodes are present in primary but not relapse and green nodes are present in the relapse but not primary. **A. AML1.** SCHISM identified a single maximum fitness 5-node tree. CPOV matrix columns and rows represent mutation clusters. Each square represents a pair of mutation clusters (I,J) and the numeric value in the square shows the fraction of mutation pairs (i,j) for which the null hypothesis was rejected (Section Vote Aggregation). **B. AML27,35,40.** For each patient, SCHISM identified a single maximum fitness 2-node tree. **C. AML15.** Two maximum fitness 3-node trees were identified. Solid arrows represent lineage relationships shared by both trees and dashed arrows represent lineage relationships specific to one of the trees. **D. AML28.** A single maximum fitness 5-node tree. **E. AML31.** A single maximum fitness 4-node tree was identified. **F. AML43.** A single maximum fitness 4-node tree. Each arrow is labeled with the fraction of maximum fitness trees that include the lineage relationship. Highlighted arrows indicate the tree manually constructed by the study authors, if it was available. GL = germline state. CL = cluster. Cluster precedence order violation (CPOV) matrices are shown to the left of each tree. Columns and rows represent mutation clusters. Each red square represents a pair of mutation clusters (I,J) for which the null hypothesis that I could be the parent of J was rejected. Each blue square represents a pair for which the null hypothesis could not be rejected.

#### Acute myeloid leukemia

This study of relapse in acute myeloid leukemia (AML) consisted of whole genome sequencing for primary and relapse samples from eight patients (AML1, AML15, AML27, AML28, AML31, AML35, AML40, AML43) [12]. For AML1, the authors identified mutation clusters with MClust [17] and manually constructed a subclonal phylogeny. For the other patients, mutation cluster means were inferred using kernel density estimation. Phylogenetic trees were constructed as in AML1 for four of the patients (AML40, AML27, AML35, AML43).

The authors described two distinct models of clonal evolution to explain relapse. In the first model, the dominant subclone present in the primary leukemia is not eliminated by therapy, but it acquires new mutations and thrives in the relapse. The patient may not have received a sufficiently aggressive treatment or may have harbored resistance mutations. In the second model, the dominant subclone is eliminated by therapy and a minor subclone in the primary acquires new mutations and thrives in the relapse, while some mutations in the primary are absent in the relapse. The mutations that allow the minor subclone to survive may have been present early on or have been acquired during or after chemotherapy, or both [12]. For patients where the authors had constructed a tree, we compared it to the best tree(s) identified by SCHISM. For the other patients, we considered whether the SCHISM trees were consistent with their suggested clonal evolution models.

For AML1, we used the published variant allele fractions and mutation cluster assignments (Table S5a in [12]). In each sample, the naive estimator was used to derive cellularity values (Supplementary Methods) for the subset of mutations which were located in diploid or hemizygous region and had coverage of ≥ 50. Hypothesis tests were performed for pairs of mutations and the CPOV matrix was constructed by voting (Methods). SCHISM identified a single maximum fitness tree, which was identical to the authors’ manually generated tree (Fig. 5A). For the remaining seven patients, we used the published cluster mean variant allele frequencies (Table S10 in [12]) and combined them with the authors’ purity estimates to infer cluster mean cellularities (Eq. S4).

Patients AML27, AML35, and AML40 were reported to harbor only two mutation clusters each, and SCHISM identified a single maximum fitness tree for each, which was identical to the authors’ tree (Fig. 5B). Patient AML15 was reported to harbor three mutation clusters, and SCHISM identified two maximum fitness trees. Each tree supported one of the authors’ two alternative models of AML relapse (Fig. 5C). In one tree, the relapse-specific mutation cluster 3 descended from the dominant subclone in the primary. This subclone consisted of the 92% of cells in the primary that carried both cluster 1 and cluster 2 mutations. In the other tree, it descended from the minor subclone in the primary (the 8% of cells in the primary carrying only cluster 1 mutations). Patient AML28 had five mutation clusters, and SCHISM identified a single maximum fitness tree. While no subclone tree for AML28 was constructed by the authors, they proposed that this patient fit the second clonal evolution model of relapse driven by a minor subclone in the primary. The SCHISM tree was consistent with this model, as the relapse-specific mutation cluster 5 descended from a minor subclone (1% of cells), which harbored only mutation cluster 1, and mutation clusters 3 and 4, which were present in the primary were absent in the relapse cluster (Fig. 5D). Patient AML31 had four mutation clusters, and SCHISM identified a single maximum fitness tree (Fig. 5E). Although no tree was provided by the authors, they proposed that this patient fit the second clonal evolution model, which was consistent with this tree. In the tree, the relapse-specific mutation cluster 4 descended from a minor subclone (21% of cells) in the primary. Cells in this subclone carried mutations in cluster 1, but not clusters 2 and 3. AML43 was reported to have four mutation clusters and SCHISM identified a single maximum fitness tree that was identical to the authors’ tree (Fig. 5F).

#### SCHISM runtime

On a MacBook Pro labptop with Intel Core i7, 4 GB of memory and 2.7 GHz CPU, a GA run with 20 generations modeling three samples and nine mutation clusters completed in 2 minutes.

## Discussion

Representing tumor evolution as a phylogenetic tree of cell subpopulations can inform critical questions regarding the temporal order of mutations driving tumor progression and the mechanisms of recurrence and metastasis. As the cost of next generation sequencing with high coverage depth decreases, many labs are employing multi-region tumor sequencing strategies to study tumor evolution. However, going from multi-region sequencing data to a subclonal phylogeny is a computationally challenging task and methods are still in their early days. Here we derived a novel framework to approach the problem. We described a statistical hypothesis test and mathematical representation of constraints on subclone phylogenies, based on rules of lineage precedence and divergence that have informed previous works in the field. We designed a new fitness function that can be used to constrain the process of subclone tree reconstruction. These tools comprise a flexible framework called SCHISM, which can be integrated with many existing methods for mutation cellularity estimation and phylogenetic reconstruction. Combined with a new implementation of genetic algorithms, we demonstrated the utility of SCHISM with simulations and by application to published multi-region sequencing studies. We were able to reconstruct the subclonal phylogenies derived by manual curation in these studies with high fidelity.

Today’s multi-region sequencing studies may often have a limited number of tumor samples, due to restrictions on the number of biopsies likely to be performed for living patients. Our results suggest that even when only a few samples are available, more accurate estimates of mutation cellularity at higher purity and coverage increase the power of the SCHISM hypothesis test. A more subtle result is that the power and Type 1 error of the test also depend on the accuracy of the standard error estimates for cellularity values. The dependency can be seen directly in the derivation of the test statistic itself (Methods Eq. 13) and indirectly in the ability of SCHISM to reconstruct complex subclone phylogenies in murine models of SCLC [13]. Although only two or three samples were sequenced from each mouse, the authors provided robust statistical estimates of mutation cluster cellularity and standard deviations.

To our knowledge, our study is the first to apply genetic algorithms to the problem of subclone tree reconstruction. Current sequencing technologies limit discovery to approximately 5-10 major subclones in patient tumor samples, equivalent to phylogenetic trees with 5-10 nodes. It is likely that in the near future, improved technology will enable discovery of a larger number of subclones. The number of topologies for a tree with *n* nodes is equal to *n*^*n*-2^ [18]. Therefore it becomes increasingly difficult to use exhaustive enumeration over all topologies when *n* = 9 (approximately 4.8 million topologies), *n* = 10 (approximately 100 million topologies), and certainly for *n* > 10. The genetic algorithm presented here enables heuristic searches over very large numbers of topologies and consequent evaluation of candidate phylogenetic trees, according to the extent to which they violated the rules of lineage precedence and divergence. However, the genetic algorithm will not always succeed when *n* is very large, and its success depends on the topology of the true tree and the distribution of mutation cluster cellularities (Supplementary Results). Alternative heuristic approaches might also prove useful in this setting such as tabu search [19, 20], simulating annealing [21, 22], or iterated local search [23].

Currently, the topology cost component of the fitness function is informed by a statistical hypothesis test that addresses a dichotomous question about the ancestral relationship of two nodes. The mass cost is a numeric measure that quantifies the extent to which the lineage divergence rule is violated, and tree fitness depends on this numeric value, rather than on acceptance or rejection of a null hypothesis. We took into consideration the power loss that would result from a statistical test based on mass cost. Such a test would depend on the estimated cellularities of a parental cluster and the sum of cellularities of its child clusters (Methods Eq. 31). A possible null hypothesis could be that the difference between parental and sum of child cluster cellularities is non-negative (no violation of lineage divergence rule), and where the absolute value of the difference measures the magnitude of the violation. This test would be underpowered compared to the topology cost test, because the expected confidence interval for the sum of mutation cluster cellularity values (Methods Eq. 2) will be larger than that for a single mutation cluster. Thus, we chose to focus our hypothesis testing framework on the topology cost. The value of our current strategy to include a numeric measure of mass cost in the fitness function is explored in (Supplementary Results and Fig. S5).

The fitness function used in this work could be further improved by incorporating measures of mutation or mutation cluster importance, using knowledge about ordering of specific driver mutations based on tumor biology, synthetic lethality, or results from single-cell sequencing. The genetic algorithm used in this work could itself be improved by the addition of online termination criteria and adaptive modulation of its key parameters, such as crossover and mutation probabilities.

A number of excellent methods to reconstruct subclonal phylogenies have been recently published [8, 7, 6, 9], and we believe all of them are likely to be useful to the cancer research community. A feature matrix comparing attributes of SCHISM to four other published methods is in Table S1. Key contributions of SCHISM are that its genetic algorithm can handle more complex tree topologies than are tractable by brute force while retaining lack of in-built bias towards linear or branched tree topologies, its modularity, and its capability to integrate information from multiple samples (biopsies from a patient) in a new statistical framework.

Finally, it is clear that under many circumstances, particularly when sample count is low and tree complexity is high, the problem of subclone tree reconstruction is underdetermined. It is likely that for at least some tumor types, the true subclone trees may be very complex. In the future, sequencing studies with a large number of samples per patient will be essential to accurately characterize these trees.

## Methods

### Framework overview

The hypothesis test and genetic algorithm used in this work are components in a general framework that decomposes the problems of mutation cellularity estimation, mutation clustering and subclone tree reconstruction (Fig. 6). Given aligned reads from whole-genome, whole-exome or targeted deep next generation sequencing, any method for mutation cellularity estimation and/or clustering can be combined with the hypothesis test described in this section. If cellularities are estimated for a cluster of mutations, the test can be applied directly to temporal ordering of clusters. If cellularities are estimated for specific mutations, the test can be applied to infer temporal ordering of mutation pairs. Given assignments of mutations to clusters, a voting aggregation scheme can be used to order the clusters themselves. The precedence order violation matrix and cluster precedence order violation matrix summarize the output of the hypothesis tests. They can be used to visualize the statistical support for potential temporal orderings (as in Figs. 3, 4 and 5). Finally, a fitness function that depends on constraints for possible values of cluster cellularities (mass cost) and the results of the hypothesis test (topology cost) can be used to rank possible topologies of subclone phylogenetic trees. The fitness function is independent of the genetic algorithm search strategy proposed in this work.

**Figure 6.**
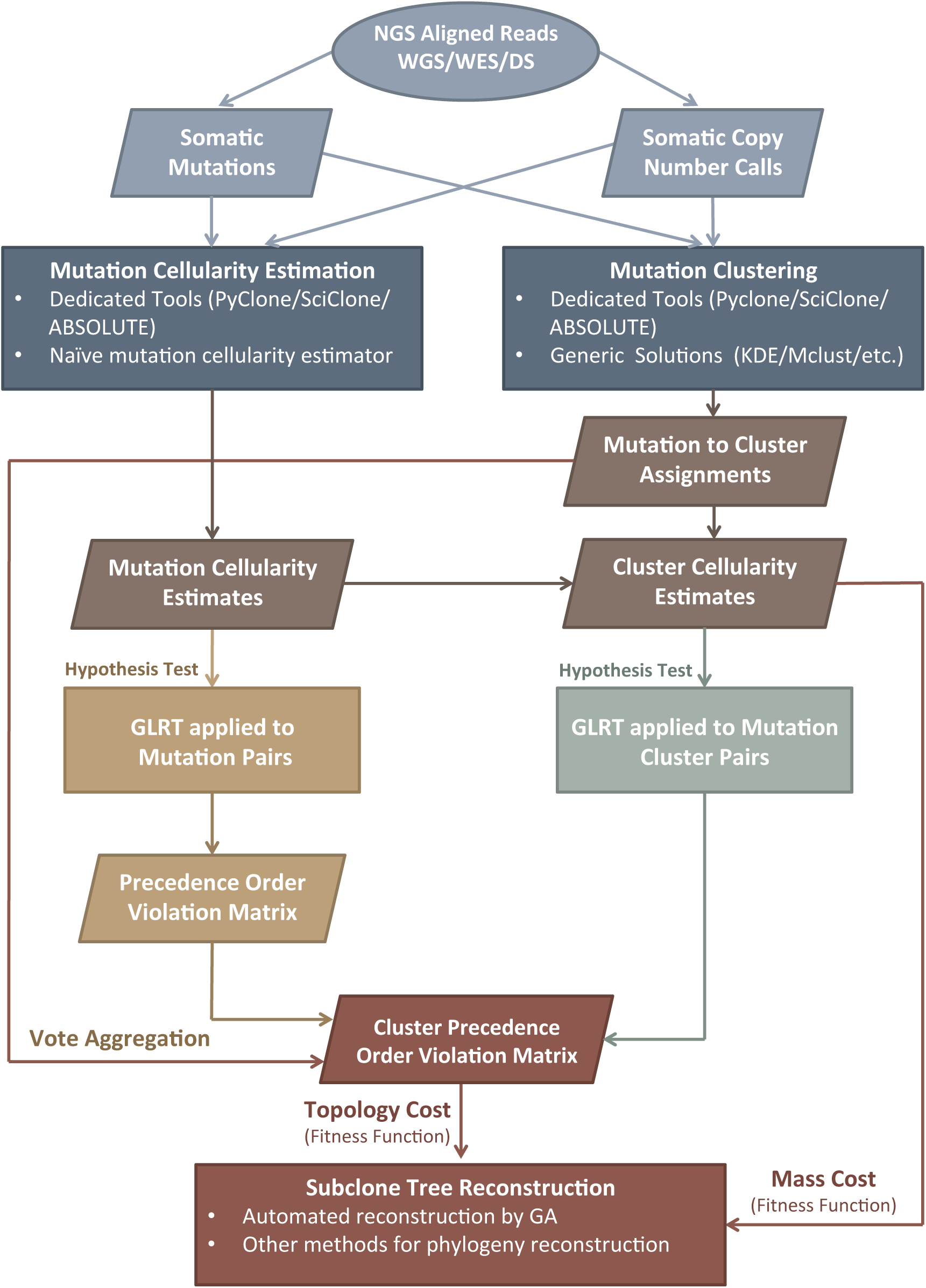
Overview of SCHISM framework. The framework decouples estimation of somatic mutation cellularities and reconstruction of subclone phylogenies. Given somatic mutation read counts from next generation sequencing data and somatic copy number calls if available, any tools for mutation cellularity estimation and mutation clustering can be applied. Their output is used to estimate the statistical support for temporal ordering of mutation or mutation cluster pairs, using a generalized likelihood ratio test (GLRT). Other approaches to tree reconstruction can be applied, by using the fitness function as the objective for optimization. GA=genetic algorithm, WGS=whole genome sequencing, WES=whole exome sequencing, DS=(targeted) deep sequencing. KDE=kernel density estimation.

### Modeling assumptions

According to the infinite sites assumption [8, 6, 7], each somatic mutation in a tumor arises only once throughout the history of the disease, and once a mutation occurs in a cell it is inherited by all descendants of the cell. It follows that given multiple tumor samples from an individual, a mutation may be present in varying proportions of the tumor cells in each sample, referred to as varying *cellularity* of the mutation across samples.

SCHISM constructs a rooted phylogenetic tree to represent the history of tumor clonal evolution in an individual. Each tree node represents cells harboring a unique compartment of mutations, defining a subclone. Each edge represents a set of mutations, acquired by the cells in the child node and differentiating them from the cells in its parental node. The somatic mutations of each tumor cell then uniquely map it to one of the nodes in the tree.

From the infinite sites assumption, we can infer that each mutation is uniquely assigned to an edge and the cells represented by a node harbor all mutations present in their parental node. Furthermore, a mutation present at a node cannot have cellularity greater than the mutations at its parental node, defining a *lineage precedence rule*. Also, the sum of mutation cellularities occuring in child nodes cannot exceed the mutation cellularity of their parent, because these mutations occur in mutually exclusive cellular populations, defining a *lineage divergence rule*.

Many methods have been proposed to estimate mutation cellularities, and our framework can be used with any of these methods. In our reconstruction of subclone phylogenies from three multi-sample sequencing studies (Results), cellularities were derived using ABSOLUTE and our own naive estimator (Supplementary Methods). Other methods such as PyClone or SciClone could also be applied.

### Hypothesis test

#### Generalized Likelihood Ratio Test

The lineage precedence rule for a pair of mutations *i* and *j*, where *i* precedes *j* in the same lineage implies that

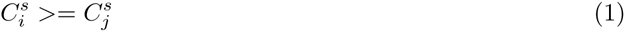

for *s* in *{*1, *…, S}*, where 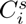 and 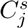 represent the cellularity of mutations *i* and *j* in sample *s* and *S* denotes the total number of samples from an individual. Then, for each ordered pair of mutations *i* and *j*, the null hypothesis 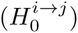 to be tested is whether mutation *i can be* an ancestor of mutation *j*, and the alternative hypothesis 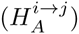 is that it is not possible for *i* to be ancestral to *j*. Let the estimated cellularity for mutation *i* in sample *s* be 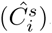. It can be represented as a draw from a normal distribution centered at the true cellularity of mutation *i* in sample 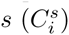 and with standard deviation 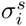.

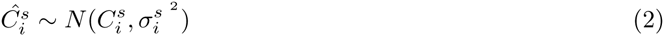

Assuming independence between the cellularity estimates for mutations *i* and *j*

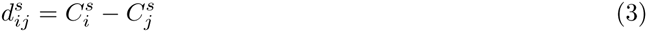

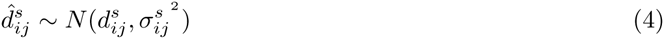

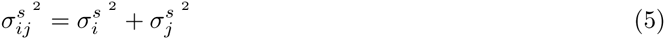

where 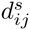 represents the true difference in cellularity of mutations *i*, and *j* in sample *s*, and 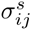is the standard deviation of 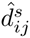, the observed difference in cellularity of mutations *i* and *j* in sample *s*. Under 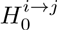, the cellularity of mutation *i* should exceed or be equal to that of mutation *j* in all tumor samples, based on the lineage precedence rule. Thus,

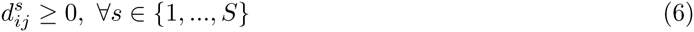

Under the alternative hypothesis 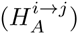, mutation *i* cannot be an ancestor to mutation *j*, and it is supported by the existence of samples in which the lineage precedence rule does not hold, and for which

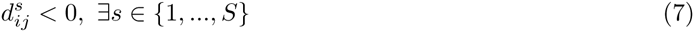

For a pair of mutations *i* and *j* and observations 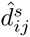 across *S* tumor samples, 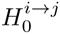 can be tested with a generalized likelihood ratio test (*GLRT*).

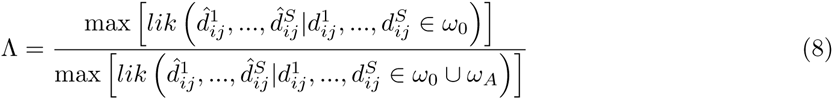

The numerator represents the maximum likelihood for observations 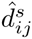 when 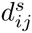 in *ω*_0_, the parameter space corresponding to 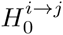 (Eq. 6). The denominator represents the maximum likelihood for observations 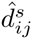 where *d*_*ij*_ is in the union of the parameter spaces for 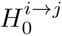 and 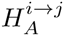, which implies there is no restriction on the values of 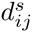.

Given the assumption of independence between 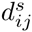 across samples and considering parameter spaces defined by 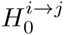 (Eq. 6) and 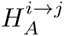 (Eq. 7), Eq. 8 can be simplified as

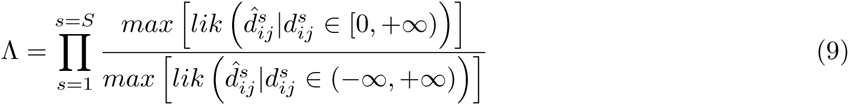

Using the normality assumption (Eq. 4), Eq. 9 is rewritten as

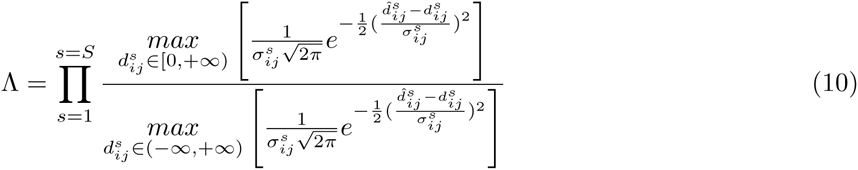

To derive the Λ, each 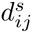 is replaced by its single sample maximum likelihood estimator under 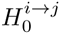 in the numerator. If 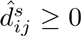, the value of 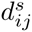 that maximizes the numerator is equal to 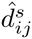. On the other hand, for 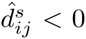, the value of 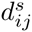 that maximizes the numerator is 0, since negative values of 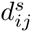 are not allowed under 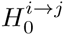. In the denominator, since the value of 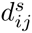 is a non-restricted parameter, its maximum likelihood estimator is always equal to 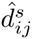. Thus Eq. 10 can be rewritten as

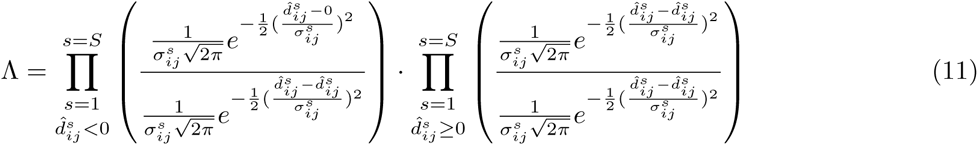

which simplifies to

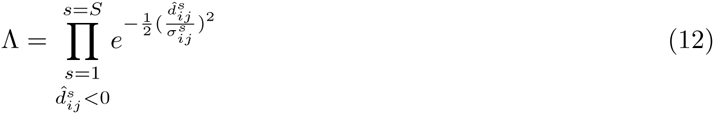

Therefore, we reject the null hypothesis that mutation *i* could be an ancestor of mutation *j* 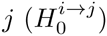 if the test statistic *T* is significantly large.

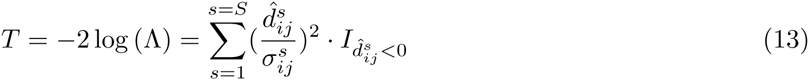

where *I* is a binary indicator variable. Eq. 13 explicitly shows how *T* depends on the accuracy of cellularity values and their standard deviation. When standard deviation is overestimated, *T* is underestimated, yielding power loss. When standard deviation is underestimated, *T* is overestimated, yielding Type 1 error inflation.

Since the standard deviation is unknown, standard error was used to calculate the test statistic.

#### Significance Evaluation

To assess the significance of an observed value of the *GLRT* test statistic (*T*) (Eq. 13), we consider the distribution of *T* under 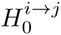 and derive the Type 1 error probability of the test. Similar results have previously been derived for a more general class of GLRTs [24]. The distribution of each summation term in Eq. 13 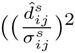.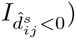 depends on the true value of 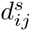 (Eq. 4). Given a fixed value of 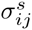, large positive values of 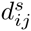 make observation of negative 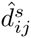 and thus a corresponding non-zero term in the summation less likely. Therefore,

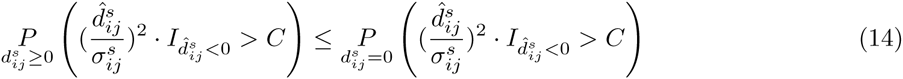

By extending the above argument to every term in the summation, we derive an upper bound for the probability of test statistic *T* exceeding a critical value *C* under the null hypothesis 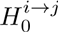.

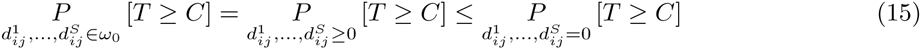

Therefore, to control the Type 1 error probability of the test, it is sufficient to control Type 1 error probability of a test where the null hypothesis is reduced to 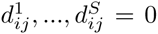 (*reduced null hypothesis*). Under the reduced null hypothesis we can derive the exact distribution of the test statistic *T* as follows.

Let 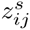denote 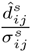

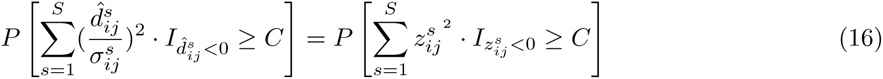

Next, let *δ*_*V*_ be a binary random variable representing the event when, for a particular subset *V* of *{*1, *…, S}*, we have 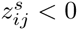 for *s* ∈ *V* and 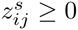 for *s* ∉ *V*. By the law of total probability,

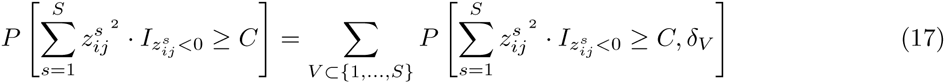

or equivalently,

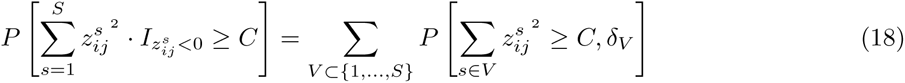

But the reduced null hypothesis (Eq. 15) states that all 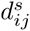 in Eq. 4 are set to zero, so

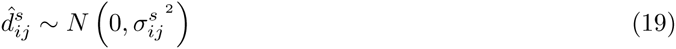

or equivalently,

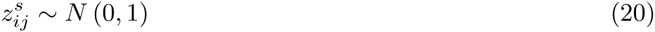

Also, for a set of independent identically distributed random draws from *N* (0, 1), the value of the summation in Eq. 18 is independent of the signs of {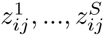}, and is a *χ*^2^ random variable with *|V|* degrees of freedom.

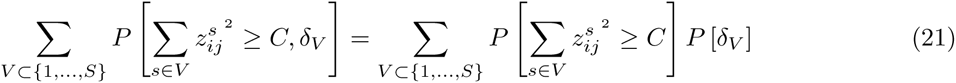

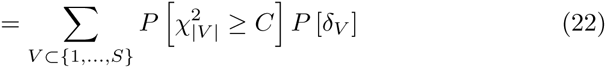

Eq. 20 implies that each random variable 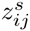 assumes positive and negative signs with equal probability. Thus, the probability of observing a particular sequence of signs for random variables 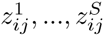, i.e. the probability of each *δ* being true, is 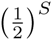.

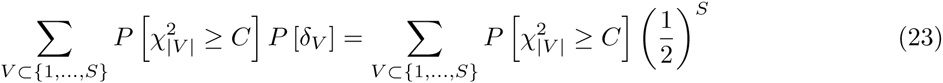

Finally, summarizing the sum above over all possible values *|V|* can take,

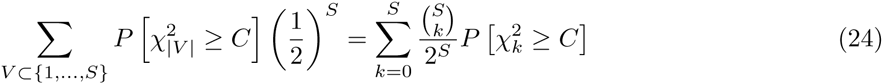

Thus it can be concluded that the distribution of *GLRT* test statistic *T* under the reduced null hypothesis is that of a random variable drawn from a mixture of χ^2^ distributions, with degrees of freedom varying in *k ∈ {*0, *…, S}*, and the weight of each mixture component equal to 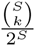. Here, a 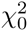 random variable is defined as one that is fixed at zero. We use this derivation to assign a conservative estimate of significance level to an observed value of the test statistic *T* under 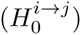.

#### Precedence Order Violation Matrix

For each possible ordered pair of mutations (*i, j*) characterized in a set of tumor samples from the same individual, we test the hypothesis that mutation *i* is a potential ancestor of mutation *j*. A fixed common significance level of α = 0.05 is assigned to decide the outcome of each pairwise test. These results can then be organized as a binary *Precedence Order Violation* (*POV*) matrix, where non-zero entries mark mutation pairs (*i, j*) for which the null hypothesis 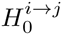 was rejected.

#### Application to mutation clusters

Given a mapping of individual mutations onto clusters, the hypothesis test can be applied to pairs of clusters rather than to pairs of mutations, and in this case is used to generate a straightforward extension of the POV matrix *Cluster Precedence Order Violation* (*CPOV* matrix) where non-zero entries mark mutation cluster pairs (*I, J*) for which the null hypothesis 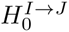 was rejected. An alternate approach for generating a CPOV matrix by vote aggregation is described next.

### Vote aggregation

The Cluster Precedence Order Violation (CPOV) matrix can be generated by the following vote aggregation approach. Let the set of mutations assigned to cluster *I* be *M*(*I*). Rows and columns of the POV matrix can be reordered so that mutations belonging to the same cluster are adjacent. Then the ordered interaction of any pair of clusters (*I, J*) is represented by a block of matrix entries with addresses

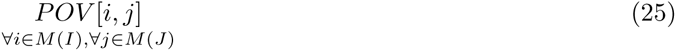

The support for potential lineage precedence of cluster *I* to cluster *J* can be summarized by a vote of the matrix elements within the block, represented as an element (*I, J*) in a cluster-level POV matrix *CPOV*

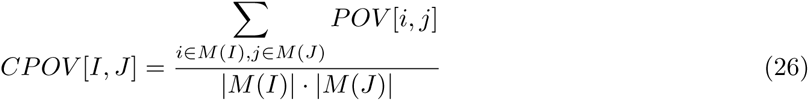

where *|M*(*X*)| denotes the number of mutations in cluster *X*.

### Genetic algorithm

A genetic algorithm (GA) is a heuristic search inspired by the process of natural selection. In an initial generation, a set of random objects is created and their fitness with respect to a fitness criteria is evaluated. Next, objects from the initial generation are selected according to their fitness to be parents of the following generation, with a preference for the fittest parents. The parental objects reproduce themselves, and their progeny may harbor new variation. The process is repeated for either a fixed number of generations or until a pre-defined convergence criteria is reached.

In our implementation, the GA searches through a space of phylogenetic tree topologies, ranking them with a fitness function that we derived based on our model assumptions. In the initial generation, we generate *G*_0_ = 1000 random tree topologies and evaluate the fitness of each tree. A sample of size 0.8 ∗ *G*_0_ trees are selected for reproduction by a fitness proportional selection method [25] and their progeny are generated, using crossover and mutation operations (Figs. 7 and 8). To increase diversity and avoid too fast convergence to a local optimum, 0.2 ∗ *G*_0_ random tree topologies are also generated. The following generation then consists of a mixture of the progeny of the previous generation(s) and new random trees, and the total number of trees is the same as in the previous generation, so that *G*_1_ = *G*_0_. The process is repeated for a fixed number γ = 20 generations. For each generation, the trees selected to be parents are not limited to the previous generation only, but can be selected from any preceding generations. To avoid getting trapped in local optima, four independent runs of the GA are performed, each with 20 generations (1000 trees per generation), and the entire ensemble of trees sampled in the four runs is ranked by tree fitness.

**Figure 7.**
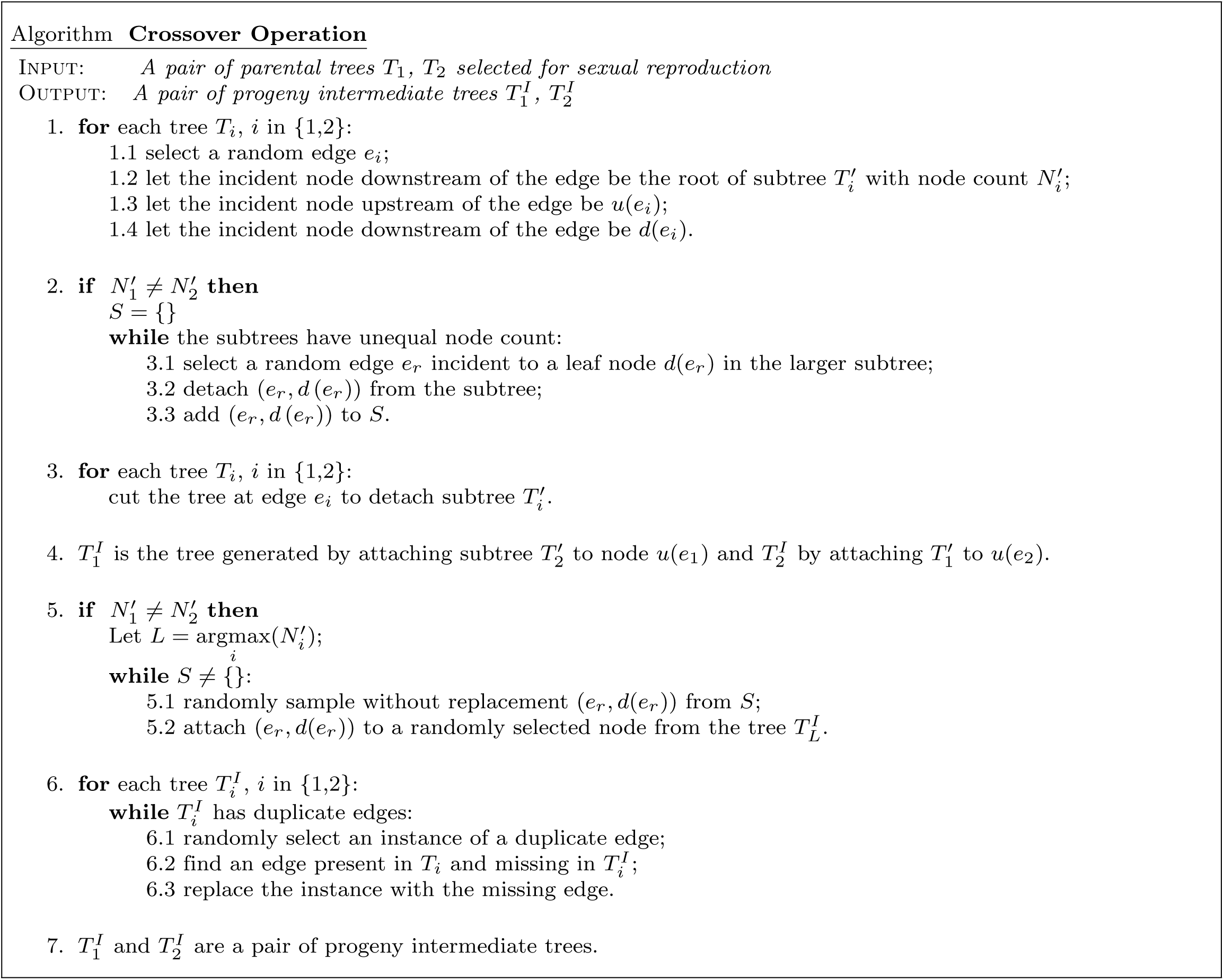
Crossover Operation. A reproductive crossover operation involving a pair of parental trees is used to generate diversity among toplogies in members of each generation produced by the genetic algorithm.

**Figure 8.**
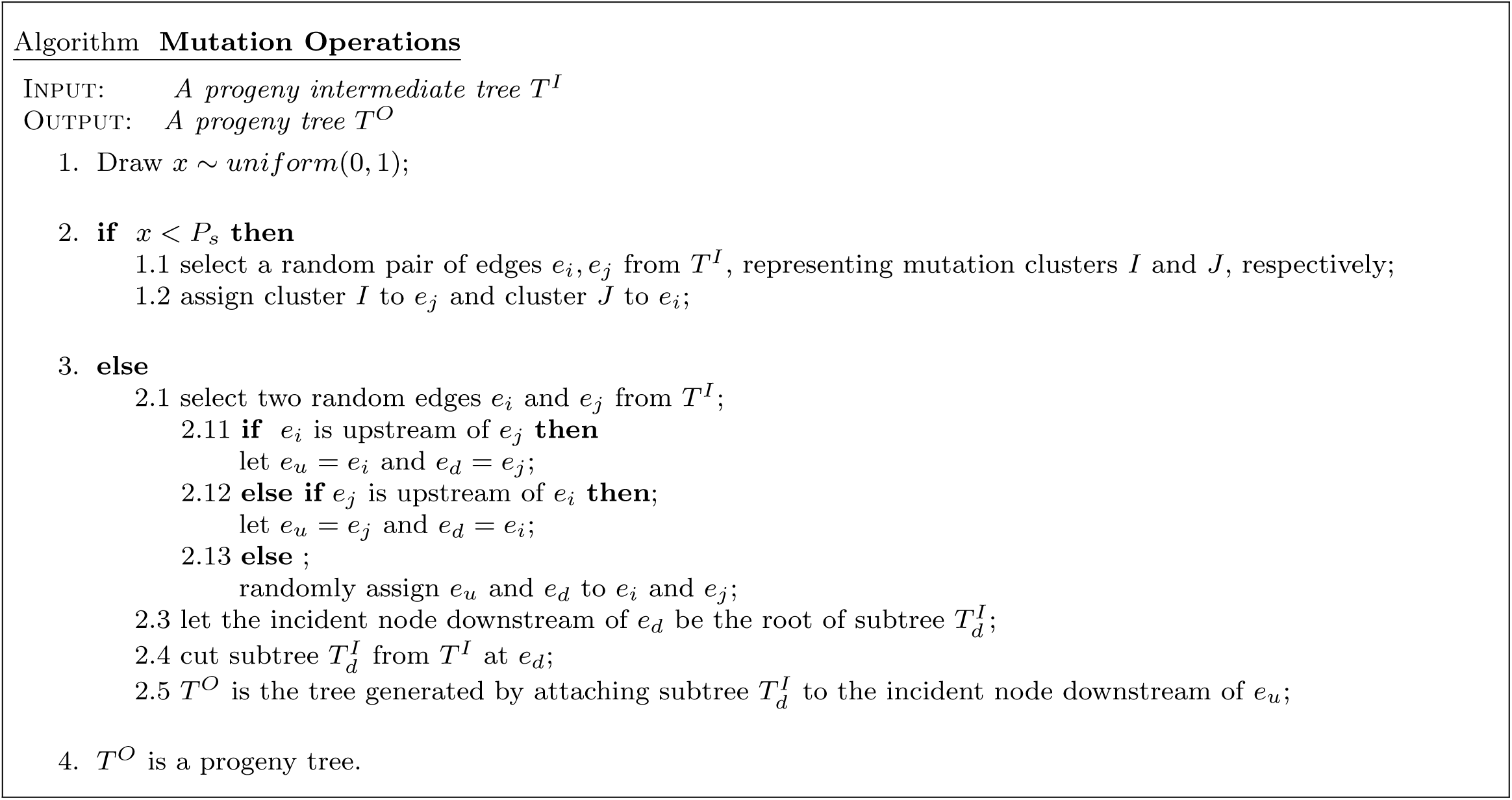
Mutation Operations. Two mutation operations are used to increase topological diversity among progeny trees in each generation produced by the genetic algorithm.

In this work, the number of mutation clusters and the cellularity of each mutation cluster in each sample is assumed to be known, and the GA is applied to explore the space of tree topologies with a given node count, including both linear and branched topologies.

#### Random Topology Generation

To generate a random tree topology, a mutation cluster is randomly selected and assigned to the incident edge downstream of the root node. An incident node downstream of the edge is appended. Next, one of the remaining mutation clusters and a non-root node are randomly selected and the cluster is assigned to the incident edge downstream of this node, again appending a new incident node downstream of the edge. The process continues until all mutation clusters in the data have been assigned.

#### Mutation and Crossover Operations

The tree topologies selected for sexual reproduction are randomly paired, and each pair undergoes a crossover operation with probability *P*_*c*_ to yield two progeny intermediate trees (Fig. 7). Next, a mutation operation is applied to each progeny intermediate tree with probability *P*_*m*_. There are two possible mutation operations (Fig. 8), and for each tree one is selected with probability *P*_*s*_. The result is a collection of new progeny trees that will appear in the next generation. (Default *P*_*c*_ = 0.25 and *P*_*m*_ = 0.9 and *P*_*s*_ = 0.6.

#### Fitness function

Tree fitness is evaluated as

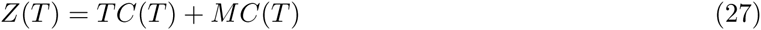

where *TC*(*T*) is a *topology cost* that summarizes violations of the lineage precedence rule, and *MC*(*T*) is a *mass cost* that summarizes violations of the lineage divergence rule.

The two components of the fitness function are useful because they represent biological properties of tumor evolution and practically, their combination can identify the true tree in cases where the topology cost or mass cost alone is insufficient (Fig. S5, Fig. S6).

The *topology cost* (*TC*) of tree *T* is

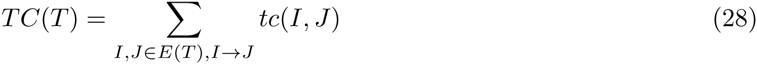

where *tc*(*I, J*) is the topology cost of each (*ancestor → descendant*) edge pair, equivalent to *CPOV* [*I, J*] (Eq. 26) and *E*(*T*) represents the set of edges in tree *T*.

The *mass cost* (*MC*) of tree *T* is

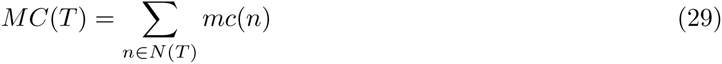

where *mc*(*n*), the total mass cost for node *n*, is the Euclidean norm of the vector of node mass costs across samples

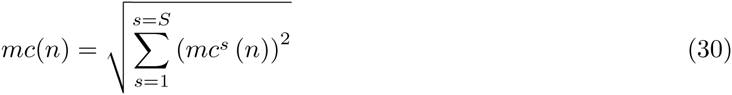

and *mc*^*s*^(*n*) is the mass cost for node *n* in sample *s*

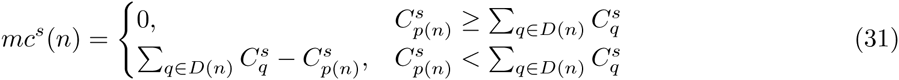

where 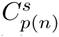 is the cellularity of the mutation cluster associated with the upstream edge incident to node *n*, *i.e.*, *p*(*n*) in sample *s*, and 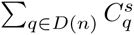 is the sum of cellularities of mutation clusters associated with its set of immediate *descendant edges D*(*n*). The fitness *F* of the tree *T* is then a monotonically decreasing function of the tree cost *Z*(*T*).

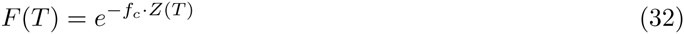

where *f*_*c*_ is a positive-valued scaling coefficient (default *f*_*c*_ = 5), yielding fitness reduction by a factor of ∼ 150X for each unit increase in total cost.

### Simulations

The simulations were designed to generate data compatible with a set of likely tree topologies and assess how well SCHISM could recover these topologies from the data. Given a tree topology, a simulation produces a set of tumor samples consistent with lineage relationships summarized in the tree. We assume that while the samples share these lineage relationships, each represents an independent instantiation of cellularity distributions over the edges of the tree. The variability among these simulated samples captures the stochastic process of preferential sampling of tumor cells in an individual’s multiple tumor samples. In each simulated sample, we model variant and reference read counts for mutations belonging to each edge in the tree, taking into account sequencing coverage depth, sample purity level and mutation cluster cellularity.

#### Generating subclonal phylogenies

Simulated trees range in size from three to eight nodes, with no restrictions on the number of child nodes. For trees with three to five nodes, an exhaustive set of topologies is generated. Otherwise, ten topologies are selected (Fig. S1). Each unique topology at a given node count is considered an *instance*. A Poisson process with rate parameter *λ* = 10 is used to simulate the number of mutations that occurred along each each edge in the tree. Node count does not include the root node, which represents the germline state prior to any somatic mutations.

A detailed description of how mutation cellularities and mutation variant allele fractions are generated in the simulations is presented in Supplementary Methods.

### Subclone size estimation

Based on our modeling assumptions, it is straightforward to conclude that in each sample *s*, the fraction of tumor cells belonging to the subclone described by node *n* in a tree can be calculated as

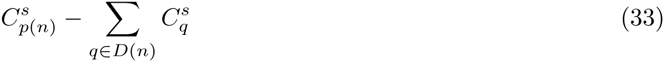

where 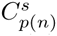 is the cellularity of the mutation cluster associated with the upstream edge incident to node *n*, *i.e.*, *p*(*n*) in sample *s*, and 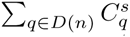 is the sum of cellularities of mutation clusters associated with its set of immediate *descendant edges D*(*n*).

### Assessment of hypothesis test

Each element of the precedence order violation matrix *POV* [*i, j*] is a binary indicator of whether the null hypothesis that mutation *i* can be ancestral to mutation *j* is rejected. Each element in the POV matrix is compared with its true value, given the correct tree topology. Performance is summarized by power and Type 1 error.

### Assessment of automated subclone tree reconstruction

The ability of the genetic algorithm to identify the correctly reconstructed subclone tree is assessed across multiple settings of key variables: tumor purity (0.5, 0.9), sequencing coverage depth (150X,1000X), tree node count (3-8), and tumor sample count (1-10). For each node count, multiple tree instances (alternate topologies for a given node count) are considered (Fig. S1). Then for each combination of settings, ten different replicates are run (Fig. S2). Each replicate can be viewed as an *in silico* patient having the selected number of samples and a distinct cellularity profile across the samples.

#### Number of maximum fitness trees identified by the genetic algorithm

In underdetermined cases, ranking of trees generated by the GA for a replicate may result in ties. We conservatively define a *Stage 1 success* for a replicate as an outcome where only one or two maximum fitness trees have been identified. To estimate the probability of Stage 1 success, the frequency of success of all tree instances and their replicates for each of 240 (2x2x6x10) unique settings of the key variables is computed. In practice, multiple maximum fitness trees can be a useful result, but limiting the number of ties in this way makes the assessment of the simulations more tractable. Note that *Stage 1 success* is only an indicator that the phylogeny reconstruction data is sufficiently determined to identify a good solution.

#### Agreement of maximum fitness tree(s) with the true tree

Even in the absence of ties, the maximum fitness tree discovered by the GA may not be the true tree that was used as the basis for the simulation. For this assessment, *Stage 2 success* for a replicate is an outcome where either a single maximum fitness or top two maximum fitness trees are the true tree. To assess Stage 2 success, we eliminate replicates where Stage 1 failed and of those remaining, calculate the fraction in which the true tree is either the maximum fitness or one of two maximally fit trees.

#### Software availability

SCHISM software is available for download at http://karchinlab.org/apps/appSchism.html

## Acknowledgments

NIH NCI grant R01CA179991 provided support for NN. We thank Christine Iacobuzio-Donahue and Alvin Makohon-Moore for valuable discussions about subclonal evolution.

## Supplementary Methods

### Naive mutation cellularity estimate

In diploid regions, a maximum likelihood estimate based on the observed variant allele fraction of a mutation can be used to infer its cellularity. We use the following simple derivation to estimate the cellularity and its standard error from reference and variant read counts for a mutation *i* in a diploid region of the genome. We further assume that the genotype of normal cells and non-variant tumor cells (tumor cells not carrying the mutation) is *AA* where *A* is the reference allele, and there is no loss of heterozygosity. Under these conditions, the probability of sampling a variant allele from tumor sample *s* with purity *α^s^*, for mutation *i* with cellularity 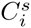 is

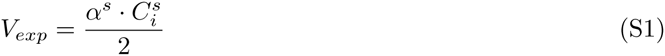

where *V*_*exp*_ is the expected variant allele fraction. By the binomial read count assumption, 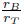 is a single sample unbiased estimator of *V*_*exp*_; here *r*_*B*_ and *r*_*T*_ represent the observed variant and total read count of a mutation, respectively.

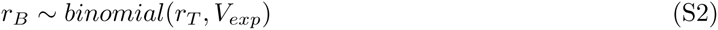

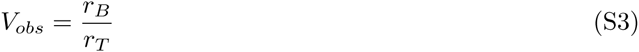

Therefore, mutation cellularity can be estimated as

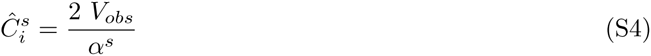

Finally, the estimated variance of the above estimator is

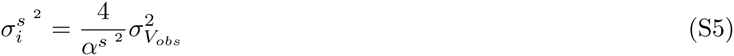

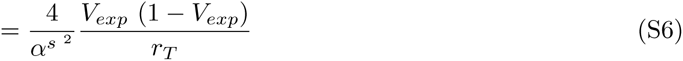

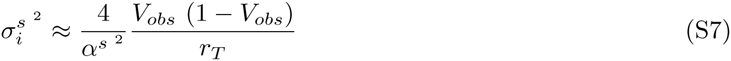

Equation S7 shows that the variance of the cellularity estimates decreases as purity and coverage increase.

Note that a simple modification allows us to extend this approach to regions with copy number = 1. For mutations in these regions, the cellularity is

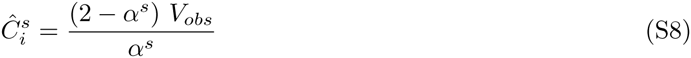

The estimated variance is

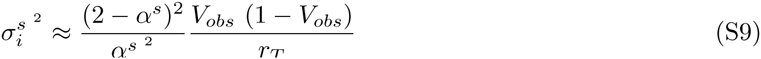

Naive cellularity estimation is useful for mutations located in diploid or hemizygous regions of cancer genomes but is of limited utility in genomes with widespread aneuploidy.

In patients with largely diploid genomes, it is possible to follow the approach above and in each tumor sample, treat the cellularity of mutations outside diploid or hemizygous regions as missing values. In the next step, for each pair of mutations/mutation clusters, the SCHISM hypothesis test can be performed—although with reduced power—by excluding any samples in which any member of the pair had missing cellularity values. On the other hand in patients with largely aneuploid genomes, more sophisticated approaches to estimate cellularity values are recommended [15, 16]. These approaches can handle variable ploidy states and by providing cellularity estimates for mutations in a larger subset of samples can help increase the power of the hypothesis test.

### Cluster cellularity estimation

By definition, all mutations in a cluster are assumed to have the same cellularity in each sample. If the cellularity of individual mutations is available, the cellularity of a cluster is estimated to be the mean cellularity of its members

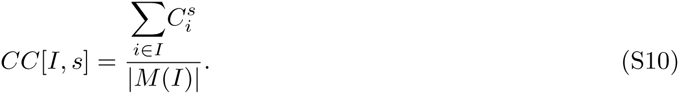

Otherwise cluster cellularity values from other sources can be used. The cellularity of each cluster across multiple samples is represented as a a matrix *CC* whose elements report the cellularity of each cluster *I*, for each sample *s*.

### Topology Similarity Measure

We apply the *Jaccard Index* to quantify the degree of similarity between two tree topologies—sharing the same mutation clusters—as follows. Let *S*_*u*_ be the set of all mutation cluster pairs (*I, J*) where mutation cluster *I* is in the same lineage as and precedes mutation cluster *J* in tree *u*. Then the similarity of two subclonal phylogeny trees *u* and *v* can be measured as:

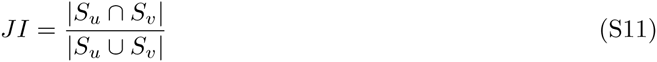

By construction, Jaccard index varies in [0, 1]. A value of 1 for the Jaccard Index indicates equality of sets *S*_*u*_ and *S*_*v*_ and thus identical topologies for trees *u* and *v*.

### Simulations

#### Generating mutation cellularities

In each simulated sample *s*, a breadth-first-search (BFS) of a subclone tree begins at the incident edge downstream of the root node. The mutation cluster corresponding to this edge has cellularity of 1, and it represents clonal mutations occurring in the most recent common progenitor cell of all the patient’s tumor cells. For subsequent edges, cellularity values are distributed with a modified version of the tree-structured stick-breaking process model [26, 6].

For each tree topology *instance* at each node count level, ten sets of mutation cluster cellularity values are generated, representing 10 samples from an individual. Each tree node then represents a unique set of cells or subclones harboring mutations, which have accumulated along the path from the root to that node. As in (Methods Eq. 31), *p*(*n*) is the mutation cluster associated with the edge immediately upstream of node *n*, and *D*(*n*) is the set of mutation clusters associated with its immediate downstream edges. Letting *n* = 0 correspond to the node immediately downstream of the root, the edge corresponding to clonal mutations is assigned cellularity 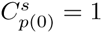. For each subsequent node *n*, the fraction of tumor cells that *p*(*n*) harbor *p*(*n*) but none of the mutation clusters downstream of *n* is 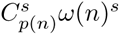 where

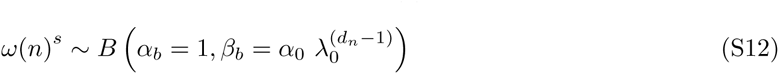

and *α*_0_ = 5, *λ*_0_ = 0.5 and *d*_*n*_ is the depth of node *n* in the rooted tree. Then 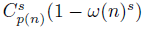 is the fraction of cancer cells harboring at least one mutation cluster in *D*(*n*) in addition to *p*(*n*). These are cells that diverge from their parental population at node *n*. Therefore,

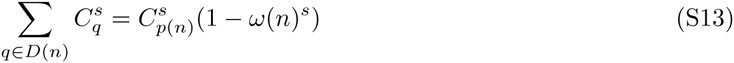

Letting *V* be a vector of size *|D*(*n*)|, and

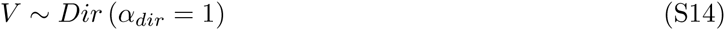

then the cellularity of each downstream mutation cluster is

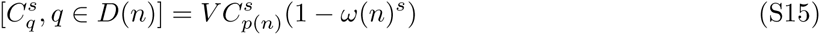

To capture variability among individuals, this procedure is replicated ten times for each tree *instance*.

#### Mutation Variant Allele Fractions

To obtain read counts that are consistent with the simulated cellularity values in each sample *s*, variant and reference read count values are generated for each mutation in mutation cluster *p*(*n*) associated with *p*(*n*) the edge immediately upstream of node *n*, given simulated cellularity value 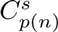, as follows. We make the simplifying assumption that mutations considered in this simulation experiment are located in diploid regions of tumor genomes in all samples. This assumption enables application of the naive cellularity estimator and allows focusing on performance of the other modules in SCHISM pipeline. Following the conditions above, the expected variant allele frequency for mutations in *p*(*n*) is

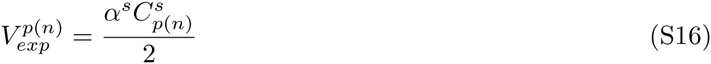

Since read counts may be overdispersed, a noisy expected variant allele frequency for each mutation *i* ∈ *p*(*n*) is generated as 

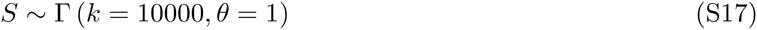

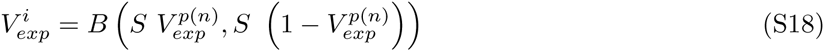

where *S* is a global precision parameter for each simulated individual. Given coverage 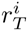 for variant *i*, variant read count is

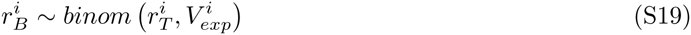

## Supplementary Results

### Genetic algorithm evaluation with large subclone count

To explore the utility of the genetic algorithm presented in this work when the number of nodes in a subclonal phylogeny is large, we designed a simulation in which 7 tumor samples were available from a patient and 15 subclones were present. In the simulations, the true subclonal phylogeny tree and the cellularity of each mutation cluster in each sample is known. Two patients were simulated. For one patient, the topology of the true tree was predominantly branched, and for the other the topology was predominantly linear (Fig. S3A, Fig. S4A). Mutation cluster cellularities for each patient are shown in (Fig. S3B, Fig. S4B). The hypothesis test was applied to each pair of mutation clusters, yielding CPOV matrices (Fig. S3C, Fig. S4C). For each simulated patient, we performed 10 runs of the GA, where each run consisted of 50 generations with 1000 trees per generation. To assess how many generations were required to identify the true tree, we monitored the progress of the GA by evaluating the topologies of all trees in the population reported to have maximum fitness after each generation. Evaluation was done by computing a Jaccard index of each of these trees with respect to the true tree (Supplementary Methods, Topology Similarity Measure). The maximum Jaccard Index after each generation is shown in (Fig. S3D, Fig. S4D).

Fig. S3 shows results for one of the simulated patients, in which the true tree topology is primarily branched. For this patient, 8 of the 10 GA runs identified the true tree at 50 generations (Fig. S3D). Fig. S3E shows the progress of one GA run that was randomly selected from these 8 runs. For each generation, the Jaccard Index for all of the maximum fitness trees is shown, rather than only for the tree with maximum Jaccard Index. At the 50th generation, the GA identified nine trees with maximum fitness and one of them was the true tree (Fig. S3E). A consensus tree of these nine trees is shown in Fig. S3F.

Fig. S4 show results for the second simulated patient, in which the true tree topology is primarily linear. For this patient, 1 of the 10 GA runs identified the true tree at 50 generations (Fig. S4D). Fig. S4E shows the progress of this GA run. At the 50th generation, the GA identified 17 trees with maximum fitness and one of them was the true tree (Fig. S4E). A consensus tree of these 17 trees is shown in Fig. S4F.

These results suggest that the GA is able to identify the true subclonal phylogeny, even in cases where the number of subclones is as high as 15, and it is reasonable to infer that it will even succeed for larger numbers of subclones, in some but not all cases. Importantly, the effectiveness of the GA may depend on details of the patient’s cancer evolutionary history and the biopsy samples.

### Limitations of topology cost or mass cost alone in identifying the true tree

Our fitness function depends on both a topology cost (TC), represented in the CPOV matrix, and a mass cost (MC). The effectiveness of the TC by itself in narrowing down candidate tree topologies and identifying the true tree depends on the topology of that true tree. For completely branched topologies, the TC alone may be sufficient in identifying the true tree, if the CPOV matrix has power of 1.0 and Type 1 error of 0. In the case of trees that include linear topologies, the TC alone will not be sufficient. Fig. S5 shows two scenarios that illustrate this point. For the 5-node branched tree (Fig. S5A) the CPOV matrix alone can identify the true tree. For the 5-node linear tree (Fig. S5B), six candidate trees (Fig. S5C) are equally likely to be the true tree if only the CPOV matrix is considered.

Fig. S6 shows an example where MC alone is not sufficient to identify the true tree, a five-node tree representing a moderately branched evolutionary pattern (Fig. S6A). Fig. S6B and Fig. S6C show cellularity values for two simulated samples and the CPOV matrix. Five candidate tree topologies (Fig. S6D) are depicted. Topogies D1, D2 and D3 have minimum TC; D4 and D5 have minimum MC. Using both TC and MC uniquely identifies the true tree topology (D1).

### Sensitivity of genetic algorithm to *f*_*c*_ parameter

The fitness function in our genetic algorithm is a decaying exponential function of tree cost. The parameter *f*_*c*_ is a coefficient of the cost and controls the rate at which changes in cost yield changes in fitness. We performed a sensitivity analysis to identify a good default value *f*_*c*_ on simulated data. Two scenarios were considered: 8-node tree with three samples and 15-node tree with seven samples. We ran 20 independent runs of the GA for each tree, using 6 unique values of *f*_*c*_. A run for the 8-node tree spanned 20 generations, and a run for the 15-node tree spanned 50 generations, to enable good sampling of tree topology space. Each generation consisted of 1000 trees. The value *f*_*c*_ = 5 produces the best results, when both scenarios are considered. For the 8-node tree, 19 out of 20 runs identified the true tree and for the 15-node tree, 17 out of 20 runs identified the true tree (Fig. S7).

### Detail of cost function for multi-sample sequencing studies

Table S2 shows the performance of SCHISM on data from the three multi-sample sequencing studies (Results) when only the topology cost (TC) or mass cost (MC) is used in the genetic algorithm. Addition of MC substantially reduces the number of maximum fitness trees identified. In this set of samples, MC alone has equivalent performance to TC+MC. As shown in (Fig. S5, Fig. S6), inclusion of both cost terms can handle scenarios where only one is insufficient.

### Comparison of SCHISM, SubcloneSeeker and TrAp output on patient AML1

Subclone Seeker and TrAp have similar inputs to SCHISM but the modeling task and outputs are different. SCHISM models a single, unified tree across multiple samples. Subclone Seeker and TrAp model trees for each individual sample and do not provide a unified tree. To illustrate these differences, we have compared the output of SCHISM to SubcloneSeeker and TrAp on eight two-sample cases (primary and relapse) from acute myeloid leukemia (AML) patients [12] and one two-sample case (primary and metastasis) from a murine small cell lung cancer (mSCLC) [13]. (Subclone Seeker can handle only up to two samples). Variant allele fractions and mutation cluster assignments from [12] were used to estimate cluster cellularities and these were input to all three methods.

Fig. S8 shows detailed results for the AML patient AML1.

SCHISM results are a unified tree for both samples in patient AML1 (Fig. S8A).

SubcloneSeeker models the evolution of each individual sample, then identifies compatible pairs of sample trees (Fig. S8B). Six trees are reported for the AML1 primary sample (Primary Tree 1, Primary Tree 6, Primary Tree 11, Primary Tree 16, Primary Tree 21, Primary Tree 26). One tree is reported for the secondary sample (Secondary Tree 1). Primary Tree 11 is reported as compatible with Secondary Tree 1 (marked with asterisks in Fig. S8B). As shown, each tree node is labeled with *n* followed by a number. The labels are not related to user-provided mutation cluster IDs. It is unclear how Primary Tree 11 and Secondary Tree 1 should be merged, given that the node numbers are disjoint between the two trees, or whether the two trees are compatible with the SCHISM unified tree.

TrAp also models the evolution of each individual sample, requiring compatibility across all samples, then outputs individual sample trees (Fig. S8C). The tree on the left represents the AML1 primary sample and the tree on the right represents the relapse sample. The TrAp trees are compatible with the SCHISM tree, which matches the manually curated tree published by Ding et al. However, the TrAp trees collapse two of the mutation clusters in the primary sample and three in the relapse sample, which is likely due to over-merging of mutation clusters.

SubcloneSeeker and TrAp appear to over-merge mutation clusters in three and two of the seven remaining AML patients, respectively. For mSCLC case mSCLC984, SubcloneSeeker does not generate any compatible pairs for the primary and secondary samples, and TrAp does not generate any trees compatible with unified trees identified by SCHISM, or the manually curated tree in [13].

---

**Figure S1.**
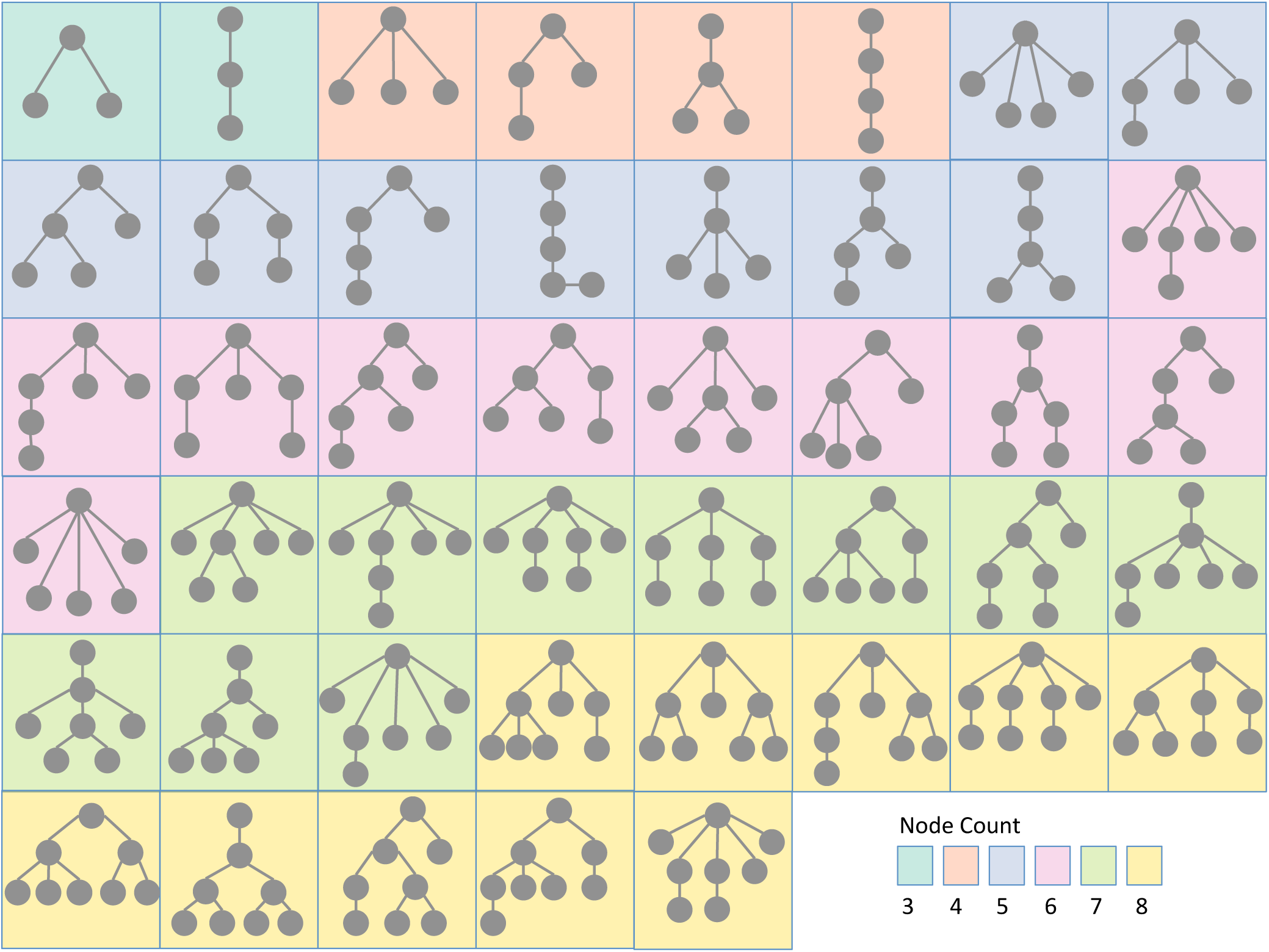
Tree topologies used in the simulations. For trees with 3-5 nodes, the exhaustive set of possible topologies were used. Otherwise, we manually selected ten topologies. Each box depicts a tree instance.

**Figure S2.**
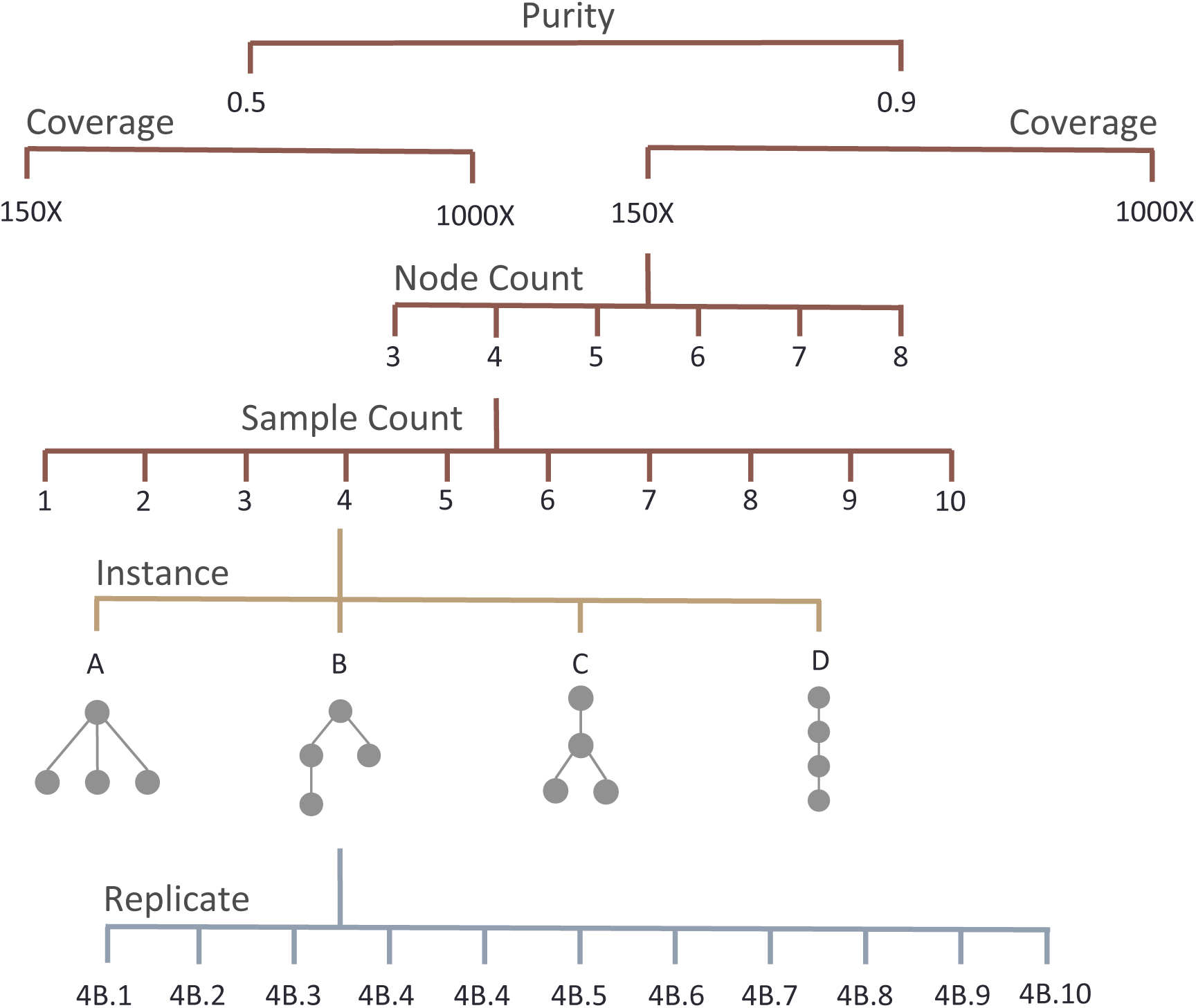
The genetic algorithm is assessed across multiple settings of tumor purity, sequencing coverage depth, tree node count, and tumor sample count. The example shows purity of 0.9, coverage of 150X, node count of 4, sample count of 4, and a selected tree instance.

**Figure S3.**
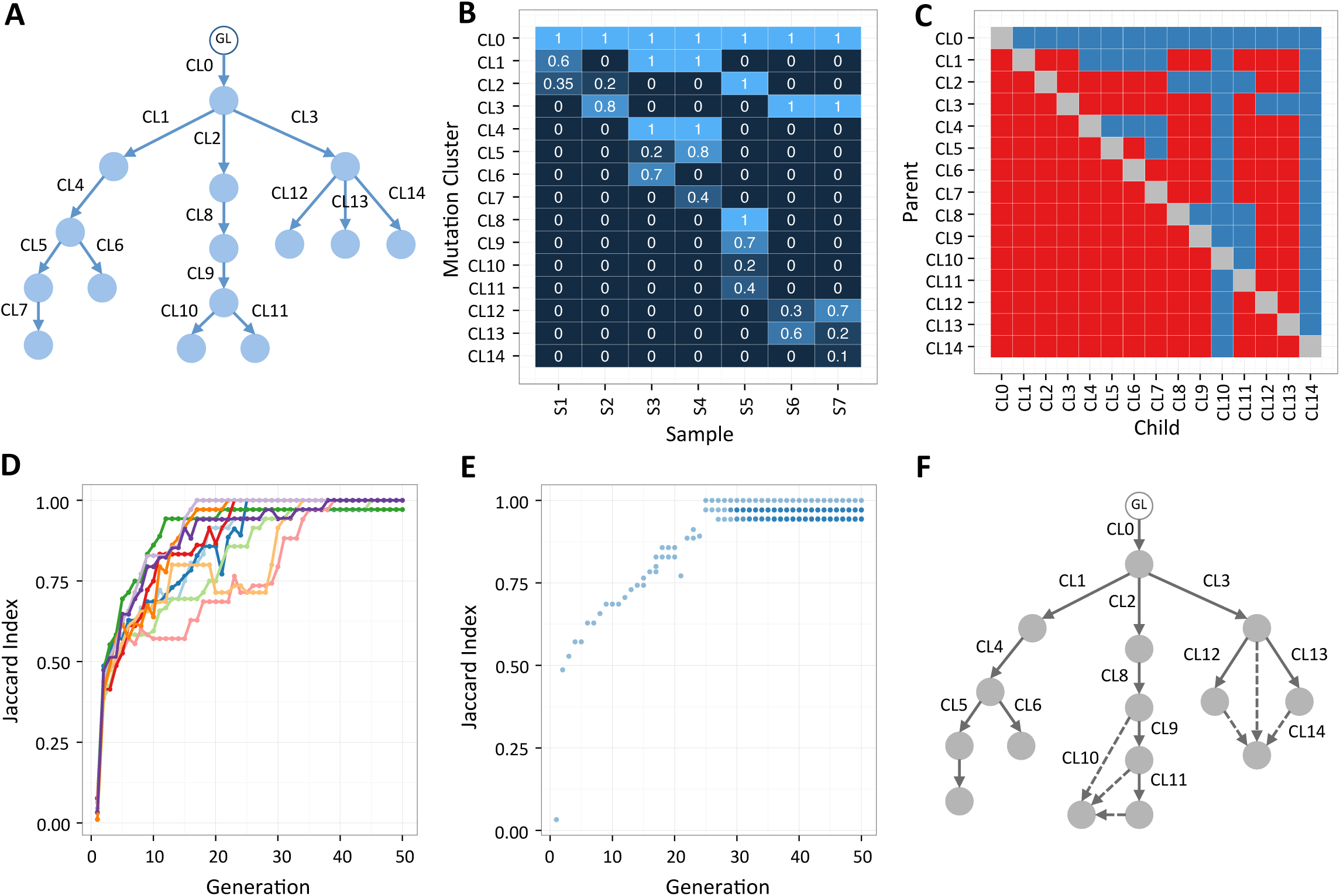
Performance of Genetic Algorithm in reconstruction of a subclonal phylogeny with large node count (n=15). **A.** a phylogenetic tree summarizing clonal evolution in a simulated patient **B.** Cellularity values of fifteen mutation clusters (CL0-14) in seven simulated biopsies S1-7 **C.** CPOV matrix depicting the hypothesis test outcomes. Each red square represents a pair of mutation clusters (I,J) for which the null hypothesis that I could be the parent of J was rejected. Each blue square represents a pair for which the null hypothesis could not be rejected. **D.** The Jaccard index (Supplementary Methods, Topology Similarity Measure) of the true tree (A) and the maximum fitness tree(s) in population at the end of each GA generation. In cases where more than a single highest maximum fitness tree was present, the maximum of the Jaccard indices is plotted. Each color trace represent one of ten independent GA runs performed on inputs in (B, C). **E.** The Jaccard index of the true tree and the maximum fitness trees at the end of each generation for a sample GA run. Each transparent circle represents a single maximum fitness tree. This GA run identified 9 maximum fitness tree topologies, one of which was the true tree (A). **F.** The consensus topology of the nine trees identified in (E). These trees shared the majority of their lineage relationships and only disagreed in parental lineage of mutation clusters CL10 and CL14.

**Figure S4.**
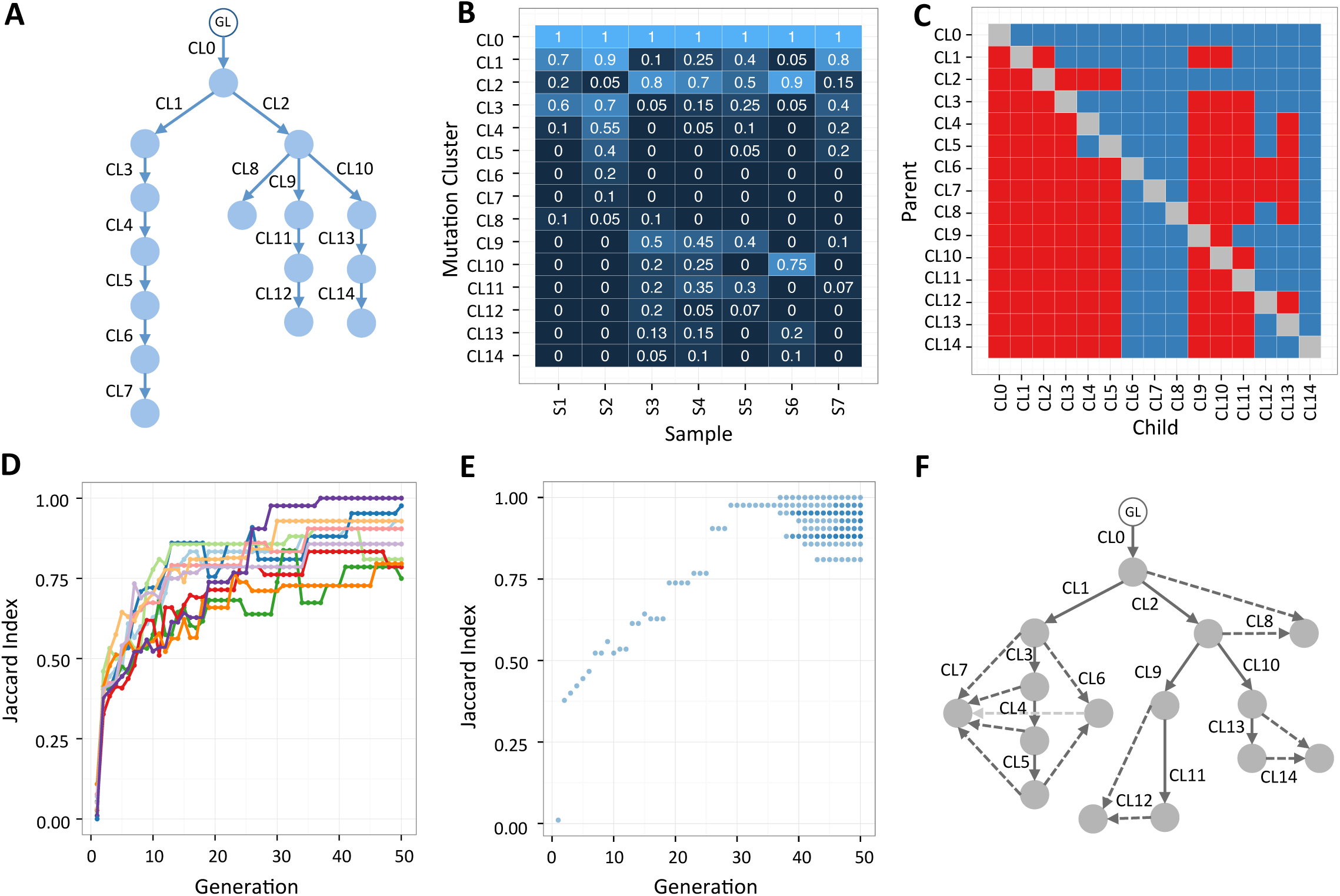
Performance of Genetic Algorithm in reconstruction of a subclonal phylogeny with large node count (n=15). **A.** a phylogenetic tree summarizing clonal evolution in a second simulated patient **B.** Cellularity values of fifteen mutation clusters (CL0-14) in seven simulated biopsies S1-7 **C.** CPOV matrix depicting the hypothesis test outcomes. Each red square represents a pair of mutation clusters (I,J) for which the null hypothesis that I could be the parent of J was rejected. Each blue square represents a pair for which the null hypothesis could not be rejected. **D.** The Jaccard index (Supplementary Methods, Topology Similarity Measure) of the true tree (A) and the maximum fitness tree(s) in population at the end of each GA generation. In cases where more than a single maximum fitness tree was present, the maximum of the Jaccard indices is plotted. Each color trace represent one of ten independent GA runs performed on inputs in (B, C). Only one of the ten GA runs (dark purple trace) identified the true tree (indicated by reaching Jaccard Index =1). **E.** The Jaccard index of the true tree and the maximum fitness trees at the end of each generation for the successful run. Each transparent circle represents a single maximum fitness tree. This GA run also found 16 other phylogenetic trees sharing the maximum fitness with the true tree.(A). **F.** The consensus topology of the seventeen trees identified in (E). These trees disagreed in parental lineage of 5 out 15 mutation clusters CL16, CL7, CL8, CL12, CL14.

**Figure S5.**
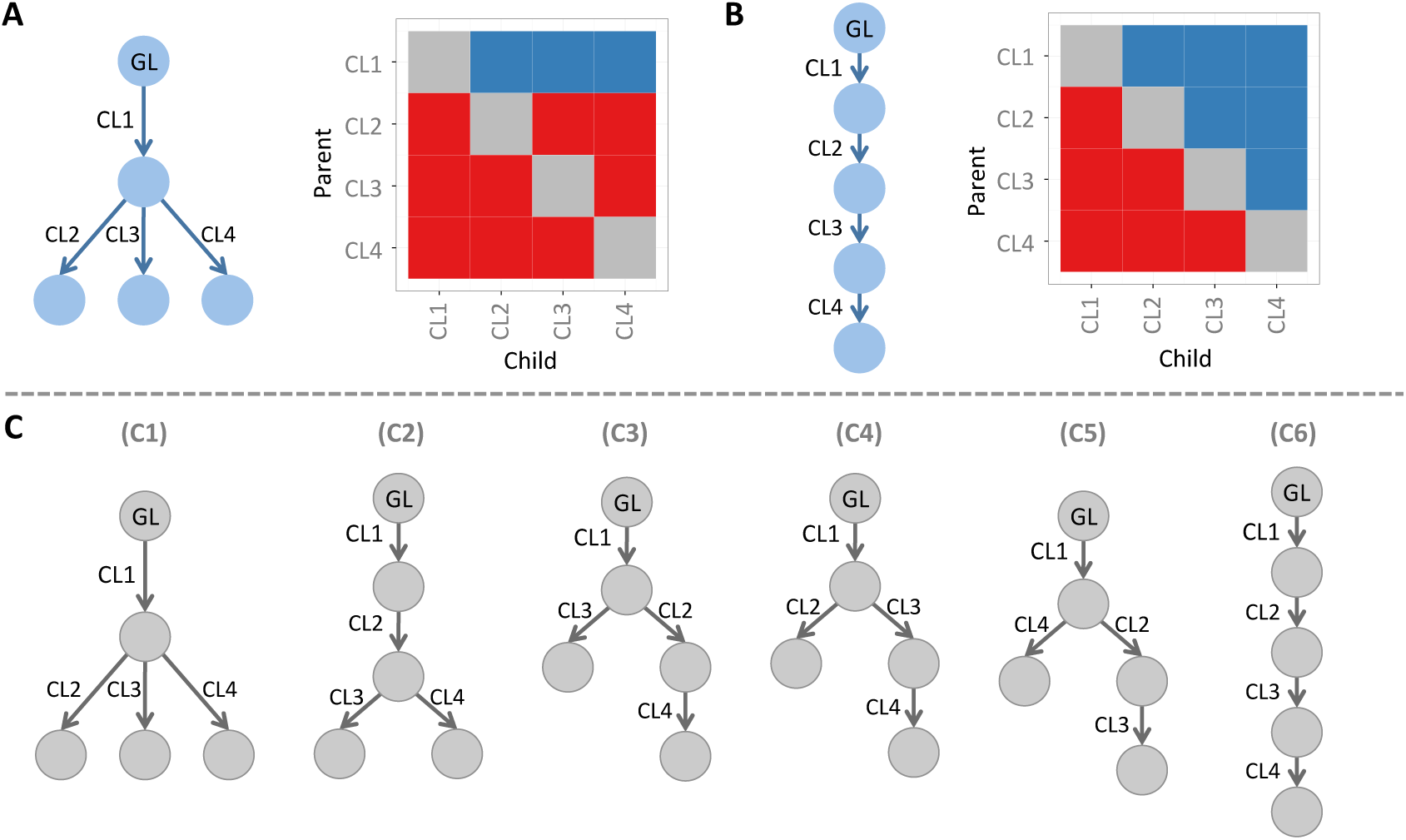
Limitations of the topology cost in identifying the true tree. Our fitness function considers both a topology cost and a mass cost. If only topology cost is used, in the case of a completely branched tree (A) and a CPOV matrix with power of 1.0 and Type 1 error of 0, the mass cost is not necessary to narrow down candidate tree topologies. However, if the tree contains linear topologies (B), the topology cost is not sufficient to identify the true tree. **A** and **B.** simple 5-node trees representing branched (A) and linear (B) evolutionary patterns. CPOV matrices under the assumption of power 1.0 and Type error 0 appear to the right of each tree. **C.** Candidate tree topologies. C1 depicts the only tree topology compatible with the CPOV matrix of the branched evolutionary pattern (A); while C1-6 are all compatible with the CPOV matrix of the linear evolutionary pattern (B), showing that in this case the topology cost alone is not sufficient.

**Figure S6.**
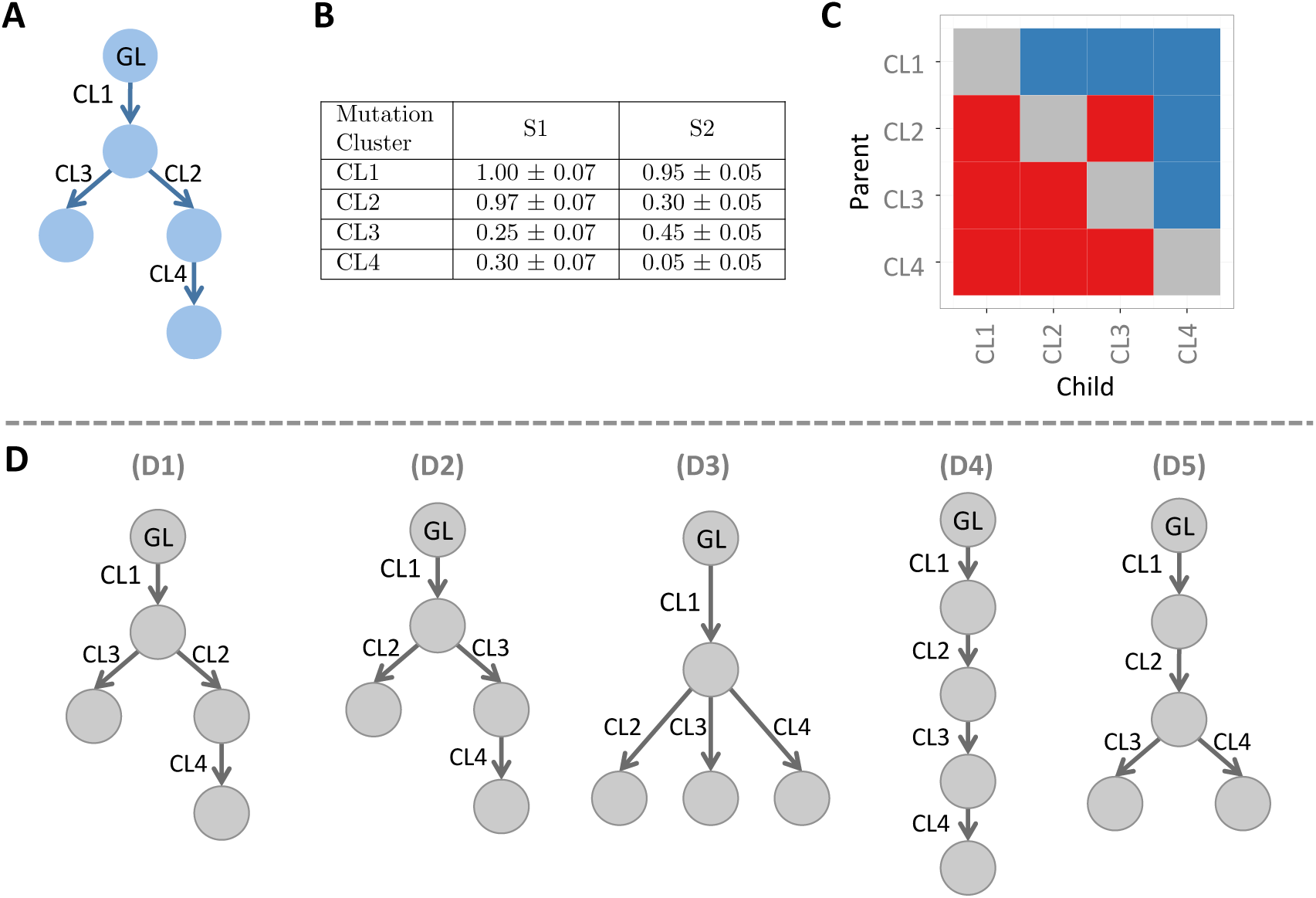
Limitation of the mass cost in identifying the true tree. Our fitness function considers both a topology cost and a mass cost. Mass cost alone is not sufficient to identify the true tree topology for the example cluster cellularity input in (B). The combination of the two cost terms outperforms each term alone in this example. **A** simple 5-node tree representing a moderately branched evolutionary pattern. **B.** Input cellularity values in two simulated samples. **C.** CPOV matrix. **D.** Candidate tree topologies. D1-3 depict topologies with minimum topology cost, and D4-5 depict those with minimum mass cost. Using both topology and mass cost uniquely identifies the true tree topology (D1).

**Figure S7.**
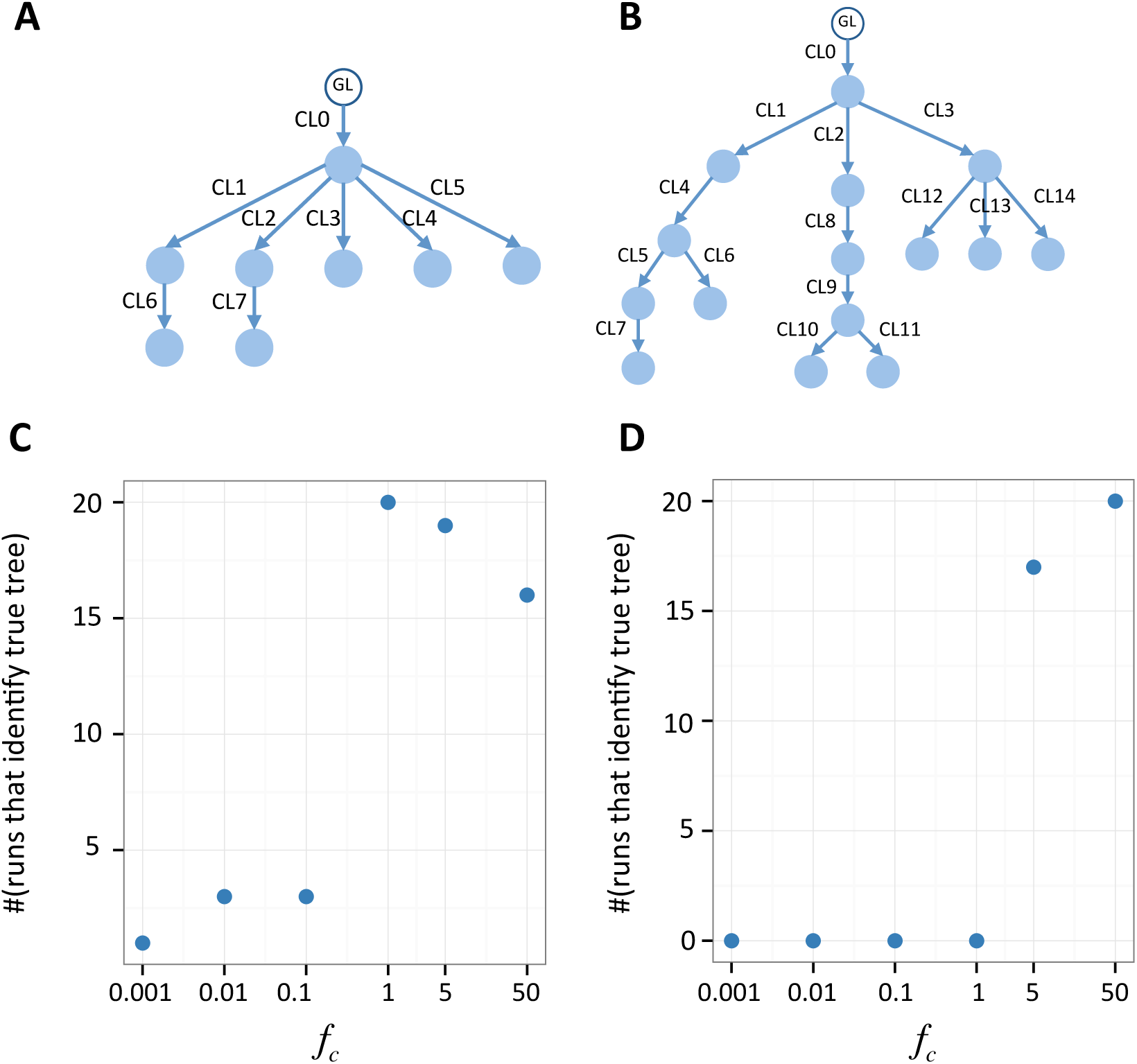
Sensitivity of genetic algorithm to *f*_*c*_ parameter. Simulations were designed to identify a reasonable default value of *f*_*c*_ in a few scenarios. An 8-node tree and a 15-node tree were simulated (as described in the Simulations section of Methods). The GA was run 20 times for each tree. For the 8-node tree, each run spanned 20 generations and for the 15-node tree, each run spanned 50 generations to enable good sampling of the tree topology space. **A** 8-node tree **B** 15-node tree. **C** and **D** The number of runs (out of 20) in which the true tree was identified by the GA for six values of *f*_*c*_. The value *f*_*c*_ = 5 produces good results for both scenarios.

**Figure S8.**
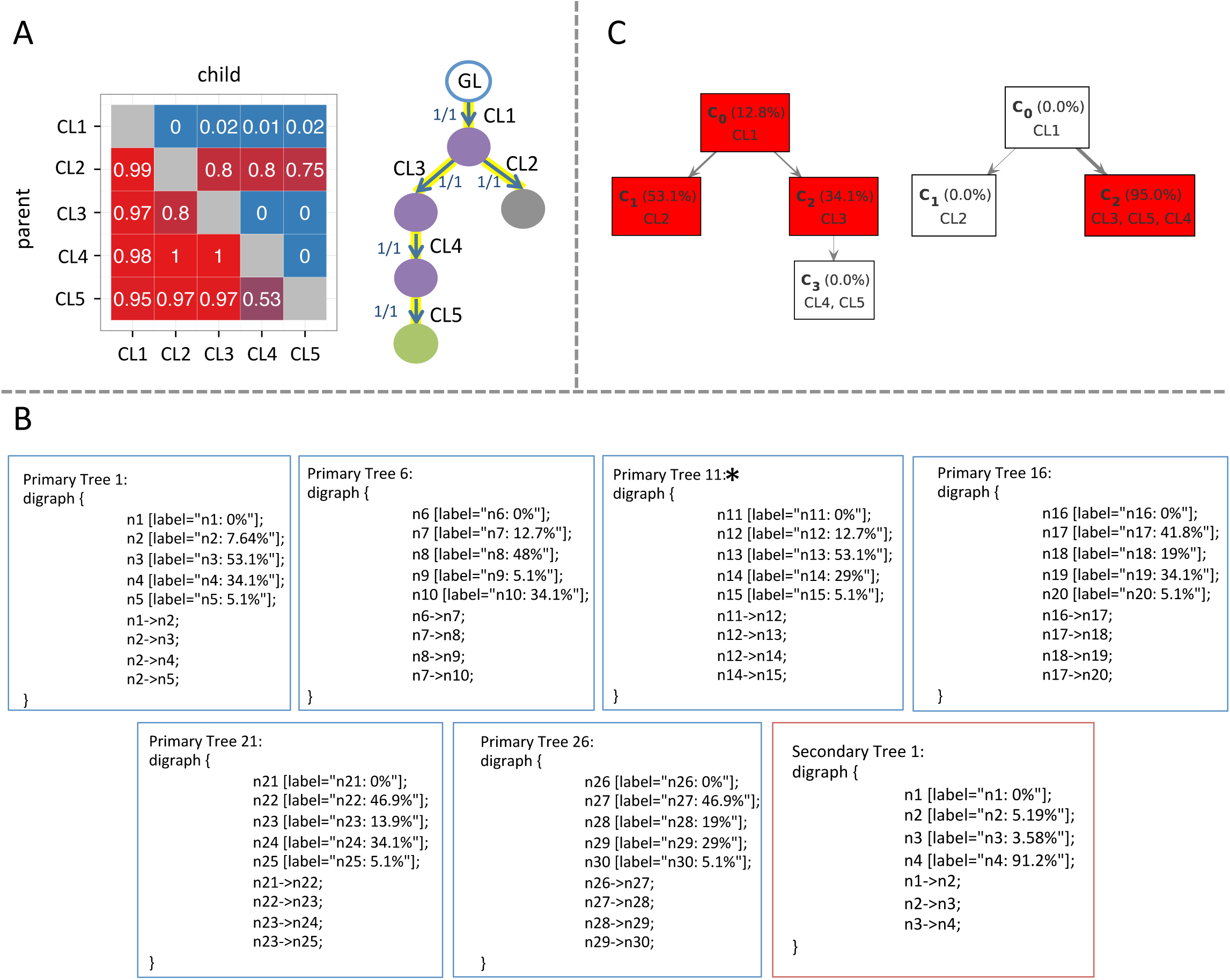
Comparison of SCHISM, SubcloneSeeker and TrAp output on patient AML1. Subclone Seeker and TrAp have similar inputs to SCHISM but the modeling task and outputs are different. **A** SCHISM models a single, unified tree across multiple samples. **B** Subclone Seeker and **C** TrAp model trees for each individual sample. The AML1 patient has one primary and one relapse sample [12]. SubcloneSeeker reports six trees for the primary sample and one tree for the relapse (secondary) sample. Primary Tree 11 is reported as compatible with Secondary Tree 1 (marked with asterisks). TrAp reports a top-scoring pair of trees – one tree (left) for the primary and one tree (right) for the relapse sample.

**Table S1.**
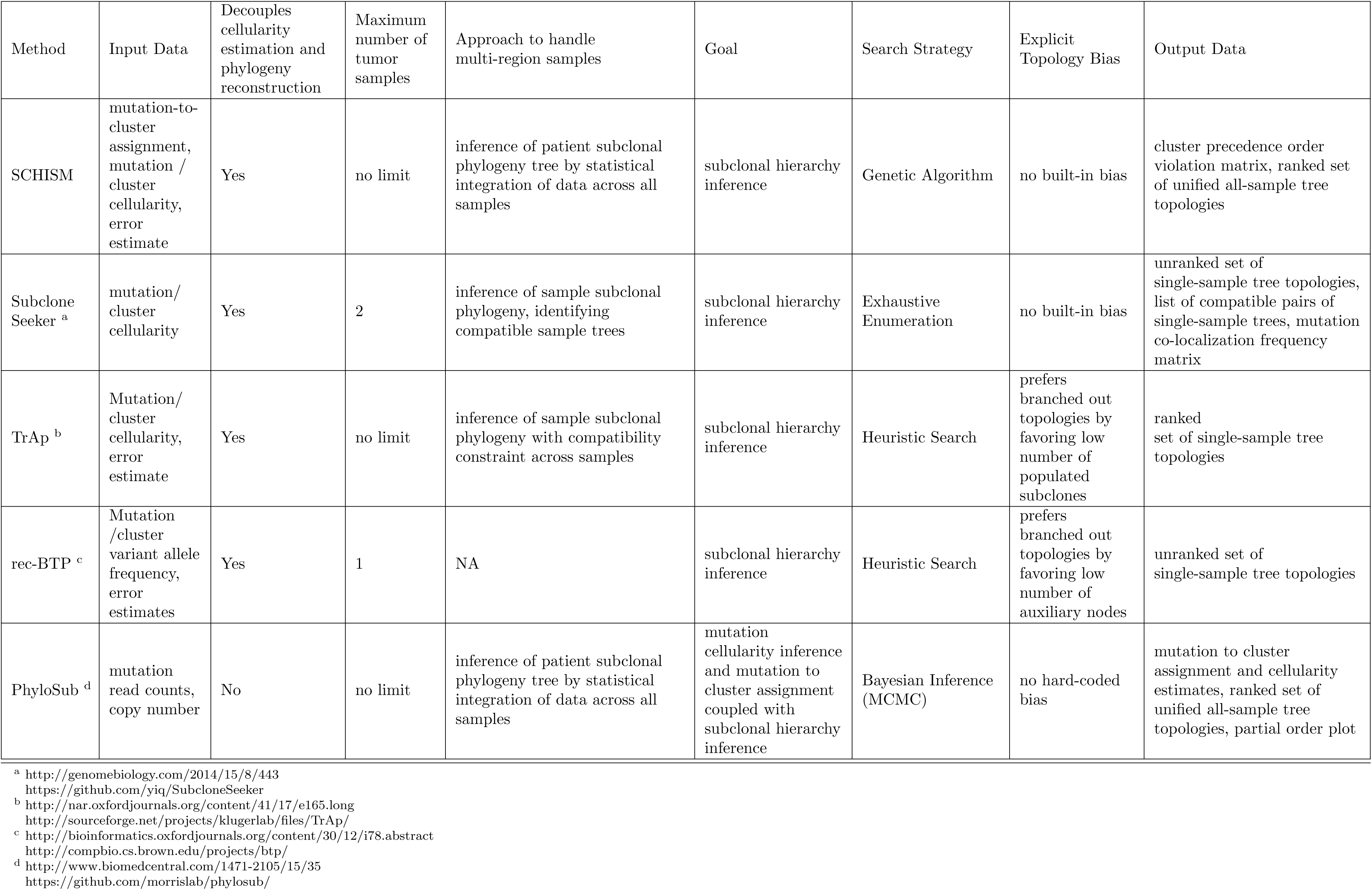
Feature matrix comparing subclonal phylogeny reconstruction methods.

**Table S2.** Detailed performance of cost function for multi-sample sequencing studies

| Sample | TC alone |  | MC alone |  | MC + TC |  |
| --- | --- | --- | --- | --- | --- | --- |
|  | #(maximum fitness trees) | includes manually crurated tree | #(maximum fitness trees) | includes manually crurated tree | #(maximum fitness trees) | includes manually crurated tree |
| AML1 | 6 | Yes | 1 | Yes | 1 | Yes |
| AML15 | 2 | Yes | 2 | Yes | 2 | Yes |
| AML27 | 1 | Yes | 1 | Yes | 1 | Yes |
| AML28 | 6 | Yes | 1 | Yes | 1 | Yes |
| AML31 | 2 | Yes | 1 | Yes | 1 | Yes |
| AML35 | 1 | Yes | 1 | Yes | 1 | Yes |
| AML40 | 1 | Yes | 1 | Yes | 1 | Yes |
| AML43 | 2 | Yes | 1 | Yes | 1 | Yes |
| CLL003 | 1 | No | 1 | Yes | 1 | Yes |
| CLL006 | 24 | Yes | 2 | Yes | 2 | Yes |
| CLL077 | 2 | Yes | 1 | Yes | 1 | Yes |
| mSCLC 3588 | 762 | No | 1 | Yes | 1 | Yes |
| mSCLC 3151 | 770 | No | 6 | Yes | 6 | Yes |
| mSCLC 984 | 24 | Yes | 6 | Yes | 6 | Yes |

